# Calpain-4 Knockdown Modulates Cholesterol Metabolism and LXRα Nuclear Localization in Alcohol-Related Liver Disease

**DOI:** 10.1101/2025.10.09.681083

**Authors:** Noriko Kitano, Jiang Li, Sam Taborski, Charis-Marie Vanderpuye, Pooja Muddasani, Sudrishti Chaudhary, Jia-Jun Liu, Silvia Liu, Juliane I Beier, Josepmaria Argemi, Ramon Bataller, Gavin E Arteel

## Abstract

**Background & Aims:** Ethanol affects lipid metabolism through multiple pathways, leading to fatty liver development in most ALD patients. Recent studies have highlighted the role of calpain, a calcium-dependent protease, in liver inflammation and fibrosis. Calpain activity is regulated by its essential subunit, Capns1, (calpain-4, Capn4), which stabilizes and modulates the activity of its catalytic isoforms, calpain-1 and calpain-2. This study investigated calpain’s impact on lipid metabolism in ALD.

**Approach & Results:** Six-week-old C57Bl6/J mice were injected with rAAV8 vectors encoding Capn4 shRNA or control vectors. After four weeks, mice underwent a 10-day period of ad libitum ethanol consumption, followed by a single gavaged ethanol administration on day 11. Following Capn4 knockdown, microvesicular steatosis was attenuated. While triglycerides and free fatty acids levels showed no significant changes, cholesterol levels were significantly reduced in the ethanol (EtOH) group with Capn4 knockdown. *Cpt1a* expression increased significantly in the EtOH group with Capn4 knockdown. Western blot analysis revealed increased Cleaved-HMGCR to Pro-HMGCR ratio in Capn4 knockdown mice, suggesting reduced HMGCR activity and suppressed cholesterol biosynthesis. LXRα expression was mainly increased in the cytoplasm in the EtOH group, and following Capn4 knockdown, it was relocalized to the nucleus via its activation. In addition, RNA sequencing analysis suggests that Capn4 knockdown contributes to the reprogramming of ethanol-induced disruptions in metabolic and homeostatic pathways, primarily those involving cholesterol metabolism.

**Conclusions:** Further investigation into the relationship between Capn4 and cholesterol biosynthesis proteins may provide insights into using calpain inhibitors as a therapeutic approach for alcohol-related hepatitis.

## Introduction

Excessive alcohol consumption is a significant global health concern, affecting millions of individuals worldwide. Chronic ethanol consumption is well known to cause dysregulation of lipid metabolism. Following heavy alcohol consumption (>60 g ethanol/day for ≥2 weeks), ∼90% of individuals develop hepatic steatosis, of whom 20–40% may develop fibrosis and 10–35% may develop steatohepatitis (1). Despite the high prevalence of alcohol-related liver disease (ALD), effective therapeutic strategies remain elusive.

Although fatty liver is often considered a benign stage of liver disease, its severity strongly correlates with progression to advanced ALD (2, 3). Early stages of ALD, including fatty liver, are typically asymptomatic and can be reversed with abstinence (4). However, due to the absence of clinical symptoms or significant abnormalities blood markers of liver damage (e.g., aminotransferases), the severity of early-stage ALD is often underestimated. Persistent alcohol intake and chronic inflammation drive progression to ASH and ultimately to advanced fibrosis and cirrhosis. While current clinical tools, including platelet counts and Fib4 index, can detect late-stage fibrosis, there is an urgent need for reliable biomarkers to identify at-risk patients during early stages of ALD (5).

Recent studies have highlighted the role of calpain, a calcium-dependent protease, in liver inflammation and fibrosis (6, 7). The activity of calpain is regulated by its essential subunit, Capns1, also known as calpain-4 (Capn4), which stabilizes and modulates its two catalytic isoforms, calpain-1 (Capn1) and calpain-2 (Capn2). Capn1 and Capn2 activation has been identified as a key driver of inflammation in other diseases of inflammation and remodeling, such as atherosclerosis and cardiac disease(8-11). Nevertheless, it’s specific role in ALD remains poorly understood. This study examined the involvement of Capn4 in ALD pathogenesis using a chronic alcohol-induced liver injury model, with particular focus on the relationships between Capn4, inflammation, and lipid metabolism.

## Method

Please see Supplemental Materials for additional details on Experimental Procedures.

### Publicly available Liver RNA Sequencing Analysis

RNA-seq data were obtained from normal livers (n = 10) and from biopsies of patients with early silent ALD (ASH, n = 11), non-severe alcohol-related steatohepatitis (AH) (n = 9), and severe AH (n = 9) from the InTEAM Consortium -– Alcohol-related Hepatitis Liver RNA-Seq study, sponsored by the National Institute of Alcohol Abuse and Alcoholism (NIAAA, USA). The study details and sequencing data can be found in the Database of Genotypes and Phenotypes (dbGAP, phs001807.v1. p1) of the National Institutes of Health (NIH, USA). The basic clinical and laboratory data of the patients included in this study, the methods used to extract RNA and perform deep RNA-seq, and the bioinformatic pipelines used to determine transcript counts have been described previously (12).

### Animals and Treatments

Six week old, male C57Bl/6J mice purchased from Jackson Laboratory (Bar Harbor, ME) were housed in a pathogen-free barrier facility accredited by the Association for Assessment and Accreditation of Laboratory Animal Care, and procedures were approved by the local Institutional Animal Care and Use Committee. Animals were allowed standard laboratory chow and water *ad libitum*. For Capn4 knockdown study, following a five-day acclimation period, the mice were divided into two groups. The first group received a 10-day \ad libitum diet of 5% liquid ethanol, with subgroups receiving either control AAV8 virus or Capn4 shRNA virus injections for 4 weeks before diet consumption (Figure 1A). The second control group received an isocaloric control diet with the same subgroups. On day 11, mice were orally gavaged with ethanol (5 g/kg body weight) or control solution. All procedures related to the feeding protocol were performed according to the chronic and binge ethanol feeding model described by Bertola et al. (13). After 9 h of ethanol administration, the mice were anesthetized with ketamine/xylazine (100/15 mg/kg, i.p.). Blood was collected from the vena cava just prior to sacrifice by exsanguination and citrated plasma was stored at -80°C for further analysis. Portions of liver tissue were frozen immediately in liquid nitrogen, while others were fixed in 10% neutral buffered formalin or embedded in frozen specimen medium (Tissue-Tek OCT compound (Sakura Finetek, Torrance, CA) for subsequent sectioning and mounting on microscope slides. The body weight of the mice and the liver weight at the time of sacrifice were measured.

**Figure 1.**
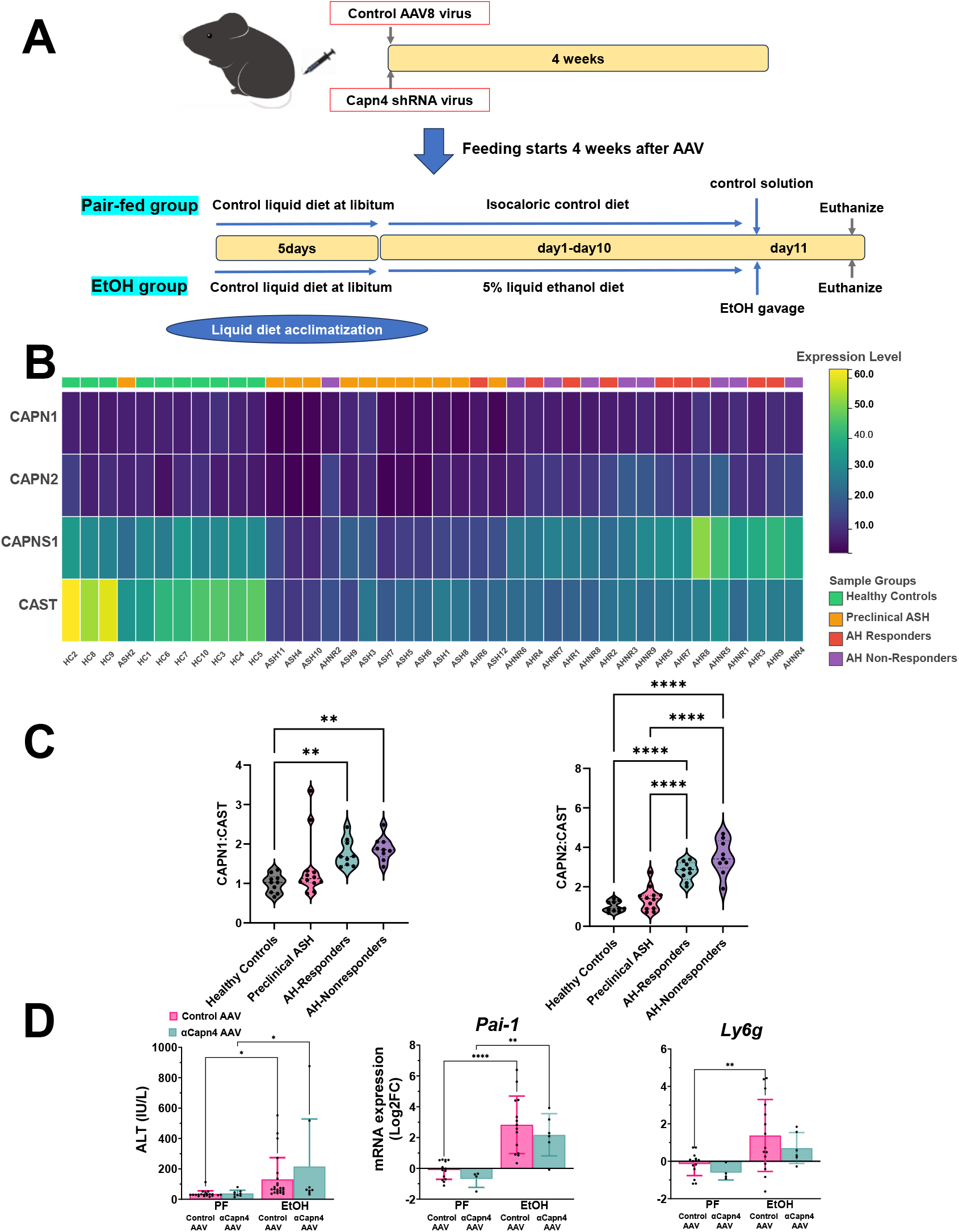
Changes in calpain family expression in human Alcohol-related liver disease and Effects of ethanol exposure and Capn4 knockdown in a mouse model. Panel A:_The schedule of the study is depicted, illustrating the protocol for ethanol administration. Mice are initially fed the control diet ad libitum for 5 d to acclimatize them to liquid diet. Afterward, ethanol-fed (EtOH) groups are allowed free access to the ethanol diet containing 5% (vol/vol) ethanol for 10 d, and control groups are pair-fed (PF) with the isocaloric control diet. <u>On</u> day 11, ETOH and PF mice are gavaged with a single dose of ethanol (5 g kg^−1^ body weight) or isocaloric maltose dextrin, respectively, and euthanized 9 h later. Panel B: RNA-seq analysis of human liver samples from healthy controls (HC), preclinical ASH (ASH), AH responders (AHR), and AH non-responders (AHNR) obtained from the publicly available dbGAP dataset (phs001807.v1.p1). This heatmap shows normalized expression levels of *CAPN1, CAPN2, CAST* and *CAPNS1* genes in each group. Panel C: Ratios of *CAPN1/CAST* and *CAPN2/CAST* were calculated from the same RNA-seq dataset as an indirect index of calpain activity. Bar graphs show the mean ± SEM of these ratios in HC, ASH, AHR and AHNR groups. Panel D To assess liver injury, we measured serum ALT level, the expression of proinflammatory genes (*Pai-1*), and immune cell markers including *Ly6g* (neutrophils). AAV, adeno-associated virus; PF, pair-fed; EtOH, ethanol.

### Biochemical assays and histology

Plasma levels of Aspartate transaminase (AST) and alanine amino transferase (ALT) were determined spectrophotometrically using standard kits (Thermo Fisher Scientific, Waltham, MA), as described previously (14). Formalin-fixed, paraffin embedded sections were cut at 5 μm and mounted on glass slides. Deparaffinized sections stained with hematoxylin and eosin (H&E) and pathology were assessed in a blinded manner. Hepatic lipids were determined using standard clinical chemistry reagents for cholesterol (CL) and triglycerides (TG) (Infinity, Thermo Fisher Scientific, Waltham, MA), for free fatty acids (FFA) (Cell Biolabs, Inc., San Diego, CA) and β-hydroxybutyrate (βHB) (Colorimetric, Cell Biolabs, Inc., San Diego, CA) (See supplemental Table1). From H&E-stained sections, lipid accumulation was quantified by steatosis scoring, where a score of 0 indicates 0% tissue coverage by lipid droplets (LDs), 1 = 1-25% coverage, 2 = 26–50% coverage, 3 = 51–75% coverage, and 4 = 76-100% coverage. Statistical analysis was performed using two-way ANOVA followed by Fisher’s Least Significant Difference (LSD) test for post hoc comparisons. For detection of hepatic Capns1 and Liver X Receptor Alpha (LXRα), paraffin-embedded liver sections were incubated with biotinylated antibodies against Capns1 and LXRα (DAKO, Carpenteria, CA). The sections were counterstained with hematoxylin (Sigma, St. Louis, MO), and immunoreactivity was visualized using a DAB detection kit (DAKO, Carpenteria, CA). All staining procedures were performed using the Vectastain kit (Torrance, CA) according to the manufacturer’s instructions. Images were captured using Metamorph software (Molecular Devices, Sunnyvale, CA).

### RNA Isolation and real-time reverse transcription polymerase chain reaction (real-time RT-PCR)

RNA extraction and real-time reverse transcription polymerase chain reaction (real-time RT-PCR) were performed as described previously (15). RNA was extracted immediately following sacrifice from fresh liver samples using RNA Stat60 (Tel-Test, Ambion, Austin, TX) and chloroform. RNA concentrations were determined spectrophotometrically and 1µg of total RNA was reverse transcribed using the QuantiTect Reverse Transcription Kit (Qiagen,Valencia, CA). Real-time RT-PCR was performed using a StepOne real time PCR system (Thermo Fisher Scientific, Grand Island, NY) using the Taqman Universal PCR Master Mix (Life Technologies, Carlsbad, CA). Primers and probes were ordered as commercially available kits (Thermo

Fisher Scientific, Grand Island, NY; see Supplemental Table 2). The comparative C_T_ method was used to determine fold differences between the target genes and an endogenous reference gene (18S). Results were reported as Log2FC of naïve control (2^-ΔΔ*C*T^).

### RNA sequencing and differential expression analysis

To investigate the transcriptomic alterations induced by ethanol administration and their modulation by Capn4 knockdown, we performed RNA-seq followed by differential expression analysis. Total RNA was extracted from liver tissues of four experimental groups: (1) pair-fed wild-type (defined as Control), (2) ethanol-fed wild-type (defined as EtOH), (3) pair-fed Capn4 knockdown (defined as Capn4 KD), and (4) ethanol-fed Capn4 knockdown (defined as EtOH+Capn4 KD) mice.

The bioinformatic pipelines used to determine transcript counts have been described previously (12). Pathway analyses were conducted using Ingenuity Pathway Analysis (IPA) software from Qiagen in Valencia, CA (http://www.ingenuity.com). This software was utilized for canonical pathway analysis and network discovery. Ingenuity pathway analysis (IPA; QIAGEN) was used to identify top canonical and enriched biological pathways determined by Differentially expressed genes (DEGs). DEGs were defined by a threshold of |log2(fold change)| > 1. Log p-value >=1.3 was selected for the study as the level of significance.

DEGs were identified using DESeq2 with p-value < 0.05 and |log_2_(fold change)| > 1. The Venn diagram was generated using the online tool *Venny* (https://bioinfogp.cnb.csic.es/tools/venny/) and *Venn Diagram Plotter* (Pacific Northwest National Laboratory) to visualize the overlap between DEGs in these comparisons.

### Immunoblots

Liver samples were homogenized in lysis buffer [20 mM Tris/Cl, pH 7.5, 150 mM NaCl, 1 mM EDTA, 1 mM EGTA, 1% (w/v) Triton X-100], containing protease and phosphatase inhibitor cocktails (Sigma, St. Louis, MO). Samples were loaded onto Invitrogen Bolt 4-12% Bis-Tris Plus gels (Thermo Fisher Scientific, Waltham, MA, USA) and subjected to electrophoresis. Gels were then blotted onto polyvinylidene difluoride membranes using the iBlot 2 PVDF Mini Stacks (Thermo Fisher Scientific, Waltham, MA, USA). The membranes were washed in TBST buffer and blocked with TBST containing 5% bovine serum albumin. Primary antibodies against HMGCR, ABCA1, ABCG1, and VCP. LXRα and GAPDH (see Supplemental Table 3) were used. After incubation with HRP-conjugated secondary antibodies, protein bands were visualized using an Enhanced Chemiluminescence kit (Pierce, Rockford, IL) and Hyperfilm (GE Healthcare, Piscataway, NJ). The results were visualized and densitometric analysis was performed using an iBright imager and onboard software from Invitrogen (Waltham, MA).

### Statistical analysis

Quantitative data are reported means ± SEM. Statistical analyses were performed using two-way ANOVA. A significance threshold of p < 0.05 was predetermined.

## Results

### Changes in calpain family expression in human Alcohol-related liver disease and Effects of ethanol exposure and Capn4 knockdown in a mouse model

Calpains are calcium-dependent cysteine proteases characterized by the calpain cysteine protease core motif in their genes (16). These enzymes regulate crucial biological processes, including cell migration, apoptosis, and synaptic plasticity through targeted protein cleavage (16, 17). The two primary isoforms, Capn1 and Capn2 are activated by calcium signaling (16, 17). Their activity is specifically regulated by calpastatin (Cast), an endogenous inhibitor that binds and colocalizes with calpain proteases (18). Capn4 functions as a small regulatory subunit that promotes and stabilizes the expression of both Capn1 and Capn2 (19-21).

To further explore the potential role of calpain family in alcohol-associated liver disease (ALD), we performed a secondary analysis using a publicly available dataset from the Database of Genotypes and Phenotypes (dbGAP, phs001807.v1.p1; National Institutes of Health, USA). RNA-seq of human liver samples from healthy controls (HC), preclinical ASH (ASH), AH responder (AHR), and AH non-responder (AHNR) revealed distinct alterations in calpain family gene expression (Figure 1B). Among these alterations, *CAPN2* expression was increased in clinical ALD (AHR and AHNR) subgroups compared with healthy controls, whereas *CAST* was markedly decreased. Since *CAST* is the key inhibitor of calpain activity, the *CAPN* to *CAST* ratio is an indirect index of calpain activity (22). The ratio of *CAPN1* or *CAPN2* to *CAST* was significantly elevated in both AHR and AHNR groups (Figure 1C), suggesting impaired calpain activity in advanced ALD.

To examine the functional significance of these observations, we established a chronic ethanol-feeding mouse model in which Capn4 was knocked down via AAV (Figure 1C). As previously observed, ethanol exposure via this regimen caused liver injury, characterized by increases in serum indices of liver damage (ALT) (23), induction of proinflammatory genes (e.g., *Pai-1*) (24) and an increased infiltration of neutrophils, as determined by hepatic expression of *Ly6g* (25, 26). These changes caused by ethanol were not significantly attenuated by Capn4 knockdown (Figure 1D). This finding suggests that Capn4 may not strongly influence acute inflammatory signaling. In addition, other markers and the expression of genes involved in liver injury were measured and Capn4 knockdown did not significantly attenuate these indices (see Supplemental Figure S1).

### Changes in calpain family expression induced by ethanol: effect of Capn4 knockdown

In pair-fed (PF) group, the expressions of *Capn1*, Capn2, *Cast* and *Capns1* (Capn4) remained unchanged by ethanol exposure at determined by RT-PCR, suggesting the mRNA expression of these genes are either minimally responsive to ethanol or regulated through alternative pathways. Following *Capn4* knockdown, the expressions of *Capn1* and *Capn2* increased in both PF and ethanol-fed (EtOH) groups (Figure 2A). In contrast, the response of *Cast* and *Capns1* varied between groups: while its expression remained stable in ethanol-treated mice following *Capn4* knockdown, PF mice showed significantly decreased *Cast* and *Capns1* after knockdown (Figure 2A). Parallel to the rtPCR results, immunohistochemical detection of Capns1 showed that EtOH exposure enhanced protein levels, which was dramatically attenuated following knockdown (Figure 2B). Taken together, these results indicate that even though transcriptional changes were modest, the αCapn4 AAV effectively prevented both basal Capn4 protein expression and the ethanol-induced increase, demonstrating successful functional knockdown at the protein level.

**Figure 2.**
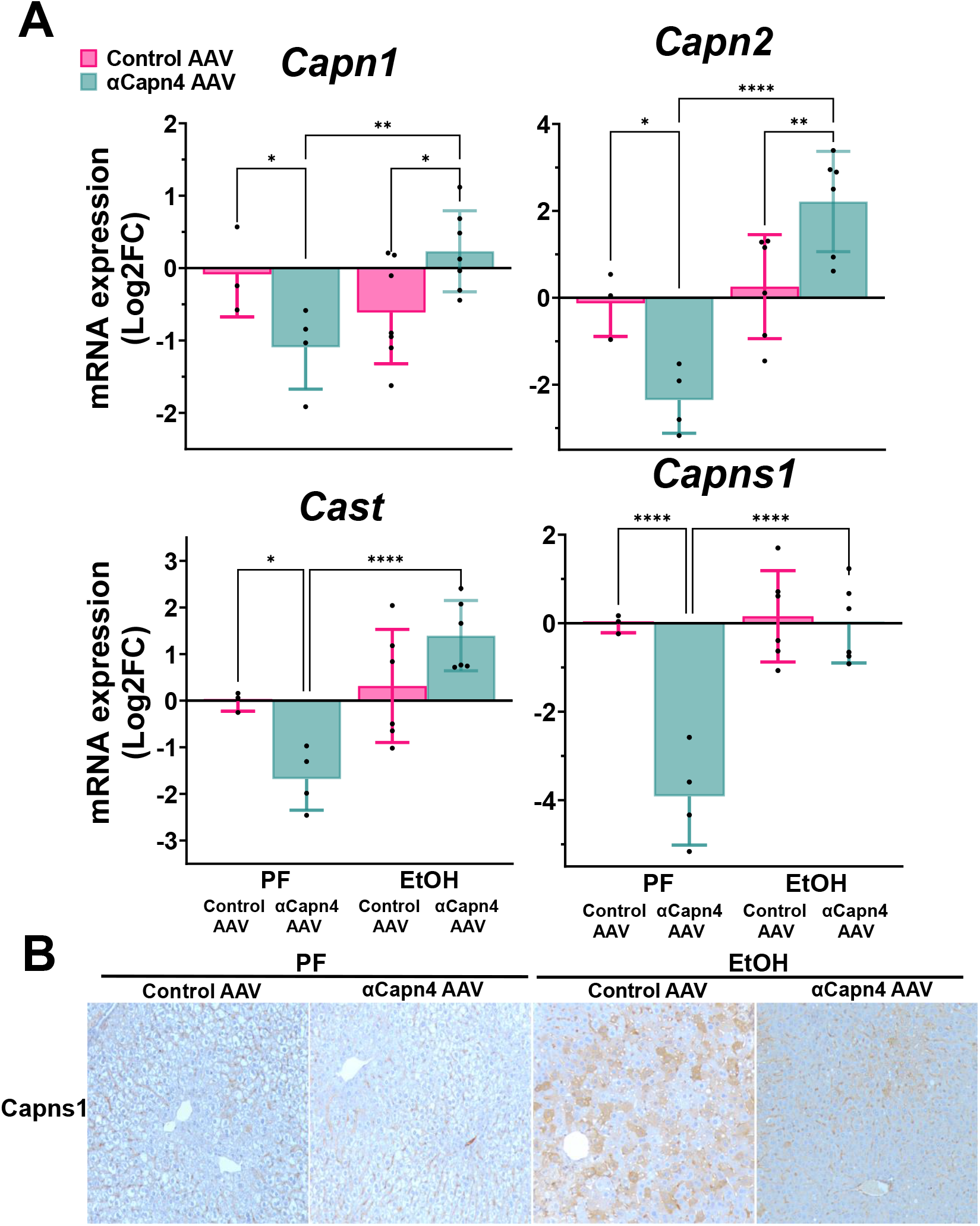
Changes in calpain family expression induced by ethanol: effect of Capn4 knockdown. Panel A: This figure illustrates changes in the expression of genes associated with the calpain system, including *Capn1, Capn2, CAST*, and *CapnS1*, between wild type and Capn4 knockdown mice. Panel B: We performed iImmunostaining for *Capns1*.

### Capn4 knockdown protected against microvesicular lipid accumulation caused by ethanol exposure

Ethanol consumption is known to cause hepatic steatosis, characterized by both macro- and micro-vesicular steatosis. Macrovesicular lipid droplets are enriched predominantly in TG (27). In contrast, microvesicular steatosis, characterized by accumulation of small lipid droplets, occurs when mitochondrial β-oxidation and/or CL metabolism is impaired (28, 29). In this study, ethanol exposure significantly increased the accumulation of microvesicular lipid droplets, which was attenuated by Capn4 knockdown (Figure 3A,C). To further characterize the specific effect of ethanol and Capn4 knockdown on lipid accumulation, total levels of TG, CL, and FFA concentrations in the liver and plasma were determined biochemically (Figure 3B). In addition, βHB in the plasma is measured (Figure 3B). Ethanol exposure caused accumulation of all three lipid pools within the liver, but only the accumulation of CL was significantly attenuated by Capn4 knockdown (Figure 3B). In addition, Capn4 knockdown significantly increased FFA and βHB in plasma in the EtOH group (Figure 3B). These results suggest that Capn4 knockdown specifically targets the microvesicular steatosis pathway by reducing hepatic cholesterol accumulation while promoting lipid mobilization and β-oxidation, as evidenced by elevated plasma free fatty acids and β-hydroxybutyrate levels.

**Figure 3.**
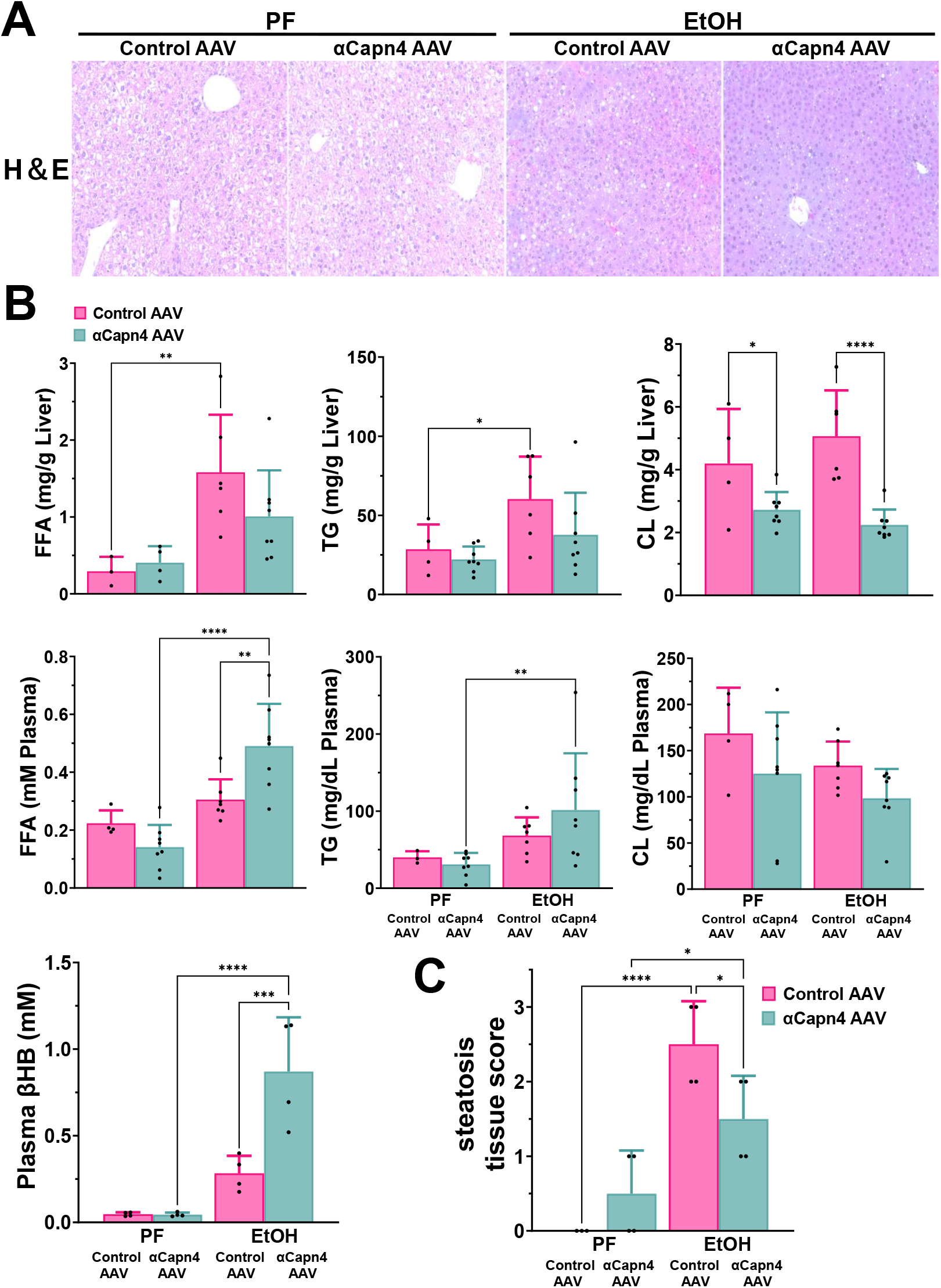
Capn4 knockdown protected against microvesicular lipid accumulation caused by ethanol exposure. Panel A: Hematoxylin and eosin (H&E; 20×) shows general histology. Panel B: Total levels of free fatty acid (FFA), triglycerides (TG), cholesterol (CL) concentrations and β-hydroxybutyrate (βHB) were determined biochemically. Panel C: Small lipid droplets were scored in HE-stained sections to evaluate the degree of microvesicular steatosis. Tissue scores were assigned based on the percentage of microvesicular steatosis observed in a single field as follows: Score 0: 0%; Score 1: 1–25%; Score 2: 26–50%; Score 3: 51–75%; Score 4: >75%.

### Capn4 Knockdown Enhances the Expression of Cholesterol-Related Genes

The expression of key lipid metabolism genes was then determined. These included genes involved in fatty acid metabolism (*Cpt1a, Fasn*), triglyceride synthesis (*Dgat2*), and CL homeostasis (*Cyp7a1, Lcat, Srebf2, Nr1h3* encoding LXRα, *Abca1*, and *Abcg1*) (Figure 4, See Supplemental Figure 2A). Ethanol exposure alone did not significantly alter the expression of any of these genes (Figure 4). However, Capn4 knockdown in ethanol-treated mice significantly increased expression of *Cpt1a* (fatty acid oxidation), *Dgat2* (triglyceride synthesis), *Srebf2* (CL biosynthesis regulation), *Nr1h3* (CL homeostasis), *Abca1* and *Abcg1* (reverse cholesterol transport) compared to the control virus group. This coordinated gene expression pattern indicates that Capn4 knockdown activates metabolic reprogramming to enhance fatty acid oxidation and cholesterol efflux.

**Figure 4.**
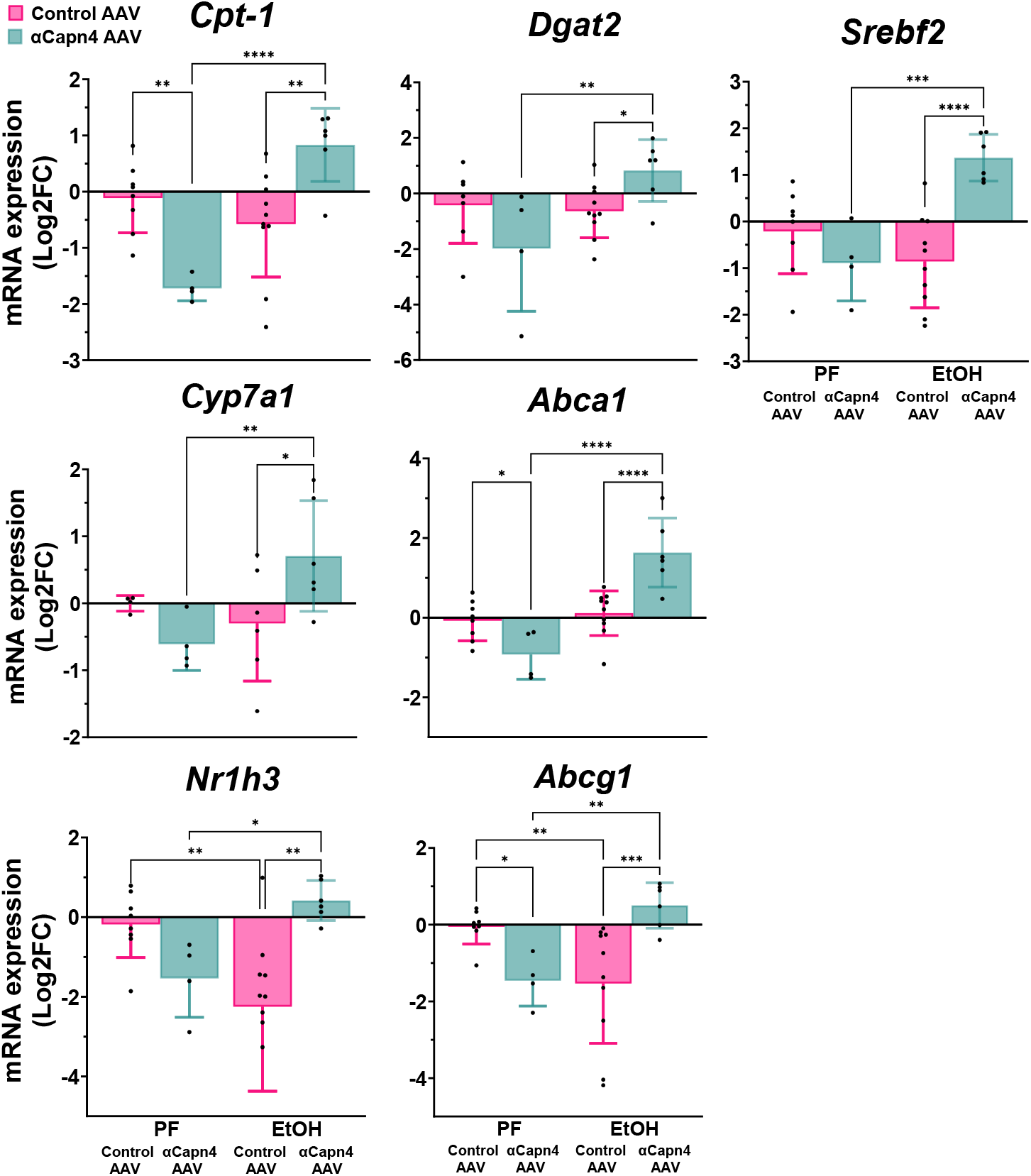
Capn4 Knockdown Enhances the Expression of Cholesterol-Related Genes. The expression of genes involved in fatty acid metabolism (*Cpt1a*), triglyceride synthesis (*Dgat2*), and CL homeostasis (*Cyp7a1, Srebf2, Nr1h3* encoding LXRα, *Abca1*, and *Abcg1*) was determined using real-time RT-PCR.

### Capn4 Knockdown Modulates Cholesterol Metabolic Proteins and Promotes Nuclear Translocation of LXRα

To further investigate the mechanism, we examined protein levels of key mediators involved in cholesterol metabolism, specifically focusing on LXRα and HMGCR expression (Figure 5). Ethanol exposure significantly altered LXRα protein expression; however, this ethanol-induced change was not affected by Capn4 knockdown. We also analyzed HMGCR, the rate-limiting enzyme in cholesterol biosynthesis, which exists in two distinct forms with different enzymatic activities. The enzymatically active form (Pro-HMGCR) can be processed by proteases to generate Cleaved-HMGCR which typically exhibits reduced or lost enzymatic activity and serves as a regulatory mechanism for HMGCR function (30, 31). Notably, Capn4 knockdown mice showed a significantly elevated ratio of cleaved/pro HMGCR, suggesting impaired cholesterol biosynthetic capacity. Additional protein expressions were also measured and are presented in Supplemental Figure 2B.

**Figure 5.**
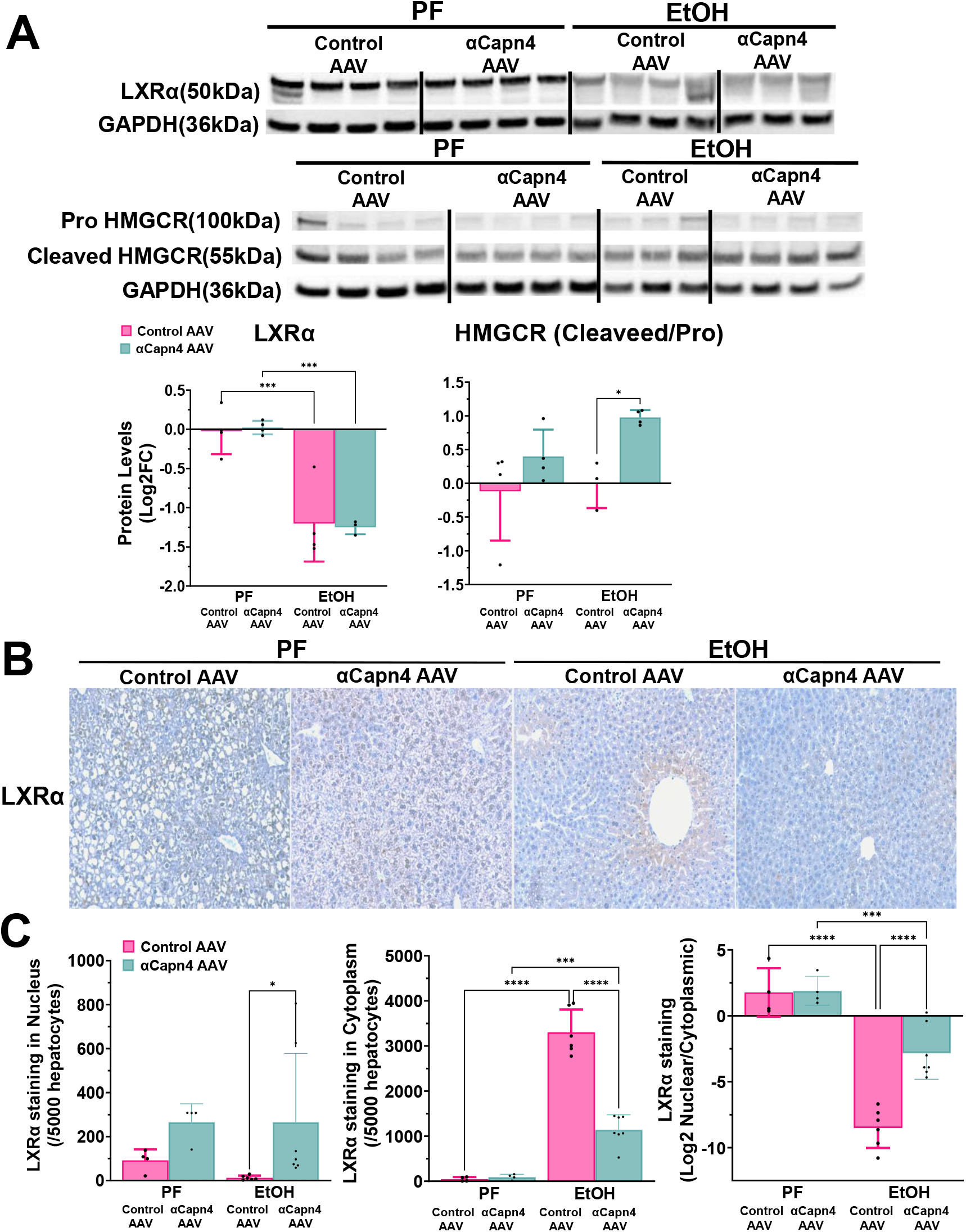
Capn4 Knockdown Modulates Cholesterol Metabolic Proteins and Promotes Nuclear Translocation of LXR_α_. Panel A: Western blot analysis was performed to evaluate the expression of HMGCR and LXRα. Additional data for ABCA1, ABCG1 and VCP are shown (Supplementary Figure S2B). Panel B: Immunohistochemical staining of LXRα in liver tissue, which indicates the nuclear and cytoplasmic localization of LXRα, is shown. Panel C: We quantified LXRα-positive nuclei and cytoplasm per 5,000 hepatocytes. Statistical analysis was performed using one-way ANOVA followed by a two-tailed Student’s t-test (*p < 0.05, **p < 0.01, ***p < 0.001).

Although ethanol exposure significantly decreased LXRα protein expression, no significant change was observed at the mRNA level (Figures 4, 5A). To further investigate the functional implications of this protein reduction, we examined LXRα subcellular localization using immunohistochemical staining. LXRα is a nuclear receptor, thereby requires nuclear localization for transcriptional activity (32). In this study, EtOH exposure increased cytoplasmic localization of LXRα in hepatocytes (Figure 5B, C). In contrast, following Capn4 knockdown, the localization of LXRα changed, with decreased cytoplasmic localization and an increased nuclear localization (Figure 5B, C).

### Transcriptomic Analysis Reveals Reprogramming of Cholesterol Metabolism by Capn4 Knockdown Under Ethanol Stress

To understand the broader transcriptional consequences of these protein-level changes, we performed RNA sequencing analysis on hepatic tissue from all treatment groups. Differential expression analysis revealed that ethanol significantly altered hepatic gene expression compared to control, as shown in the volcano plot (Figure 6A, left panel). Capn4 knockdown under ethanol exposure induced a distinct set of transcriptional changes compared to ethanol treatment alone (Figure 6A, right panel), suggesting that Capn4 modulates the transcriptomic response to ethanol in a calpain-dependent manner. Venn diagram analysis demonstrated modest overlap (39.2%) between differentially expressed genes from the ethanol + Capn4 knockdown versus ethanol and ethanol versus control comparisons (Figure 6B), with 30.8% and 30% of genes being unique to each comparison, respectively. This pattern indicates that Capn4 knockdown substantially reprograms, rather than simply reverses, ethanol-induced transcriptional changes. Pathway analysis using Ingenuity Pathway Analysis (IPA) revealed striking differences between treatment groups (Figure 6C). While ethanol exposure robustly activated biosynthetic pathways, most notably cholesterol biosynthesis (Z-score >4), compared to control, Capn4 knockdown under ethanol exposure suppressed these same cholesterol biosynthetic pathways (Z-score <0). Additional pathways affected included LXR/RXR activation, which was prominently upregulated by ethanol but showed different regulation patterns following Capn4 knockdown. These transcriptomic findings directly complement our protein-level observations of LXRα relocalization and suggest that Capn4-mediated calpain activity coordinates both LXRα subcellular localization and downstream cholesterol metabolism gene expression programs in response to ethanol exposure. Detailed results of pathway and upstream regulator analyses are provided in Supplemental Table 4.

**Figure 6.**
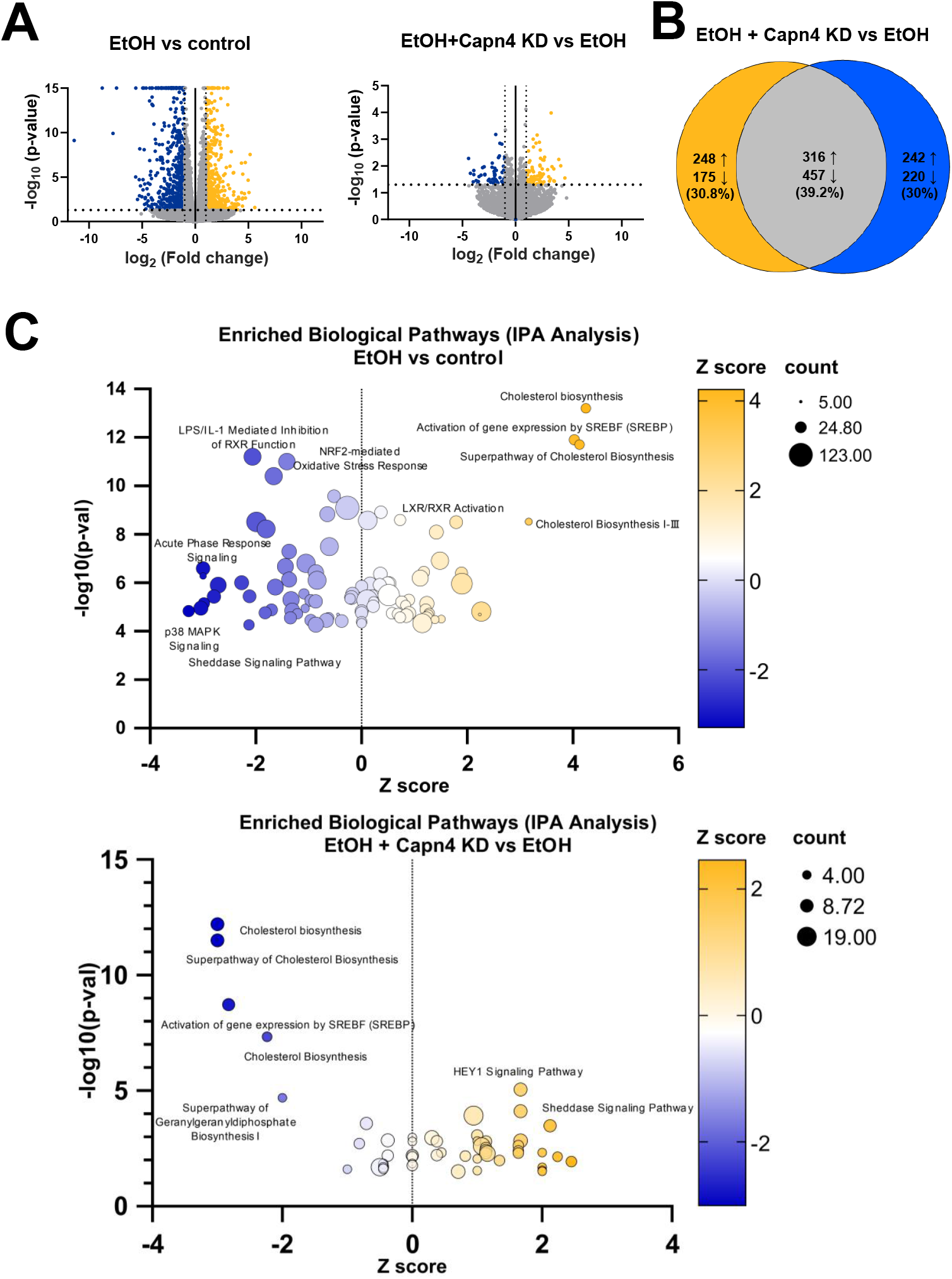
Transcriptomic Analysis Reveals Reprogramming of Cholesterol Metabolism by Capn4 Knockdown Under Ethanol Stress. Panel A: Volcano plots showing differentially expressed genes (DEGs) in EtOH vs Control (left) and EtOH + Capn4 KD vs EtOH (right). Cutoffs: |log_2_FC| > 1 and adjusted p-value < 0.05. Panel B: Venn diagram illustrating the overlap of DEGs between EtOH + Capn4 knockdown and EtOH. Panel C: Canonical pathway analysis (IPA) of DEGs in EtOH vs Control and EtOH + Capn4 KD vs EtOH. Among the top 100 canonical pathways ranked by p-value, only those for which an activation z-score could be calculated are presented in the pathway chart. IPA, Ingenuity Pathway Analysis; KD, knockdown.

## Discussion

In this study, we investigated the role of Capn4 knockdown in lipid metabolism in ethanol-treated mice. Ethanol is known to promote hepatic lipogenesis and concurrently suppress β-oxidation (33). However, the impact of ethanol on CL metabolism remains inconclusive (25, 33). This findings demonstrate that Capn4 knockdown selectively attenuates cholesterol accumulation without affecting TG or FFA levels (Figure 3). This suggests a more pronounced involvement of calpain in CL metabolism within the context of alcohol-related liver disease (ALD). Additionally, Capn4 knockdown markedly attenuated ethanol-induced microvesicular steatosis (Figure 3), indicating that changes in CL metabolism may contribute to the amelioration of lipid droplet accumulation in the liver.

RNA-seq analysis revealed that Capn4 knockdown suppressed the ethanol-induced activation of CL biosynthetic pathways in ethanol-treated mice. Capn4 encodes the essential small regulatory subunit shared by classical calpains (calpain-1 and -2), and its deletion results in embryonic lethality before day E11.5 in mice, underscoring its indispensable regulatory function (34). In the present study, Capn4 knockdown alone also exerted marked effects on pathways related to anabolic responses and broader metabolic regulation, consistent with a role as a master regulator (see Supplemental Table 4). Collectively, these findings support the notion that, in response to ethanol, Capn4 knockdown reprograms hepatic gene expression and plays an important role in maintaining hepatic CL homeostasis during alcohol-induced liver injury.

We observed increased expression levels of genes involved in CL homeostasis, including *Serbf2, Cyp7a1, Nr1h3* (encoding LXRα), *Abca1*, and *Abcg1*, in Capn4 knockdown mice (Figure 4). This upregulation is likely a response to decreased intracellular CL levels, leading to the increase in transcription of *Srebf2* and *Nr1h3* (35, 36). SREBP2 promotes CL biosynthesis via HMGCR and uptake via LDLR (37, 38). Interestingly, despite increased Srebf2 expression in Capn4 knockdown mice, HMGCR activity was decreased. Previous studies have shown that Capn2 is involved in HMGCR degradation, suggesting that HMGCR is a likely substrate of calpains (39). HMGCR degradation is primarily regulated by the ubiquitin-proteasome system in response to CL levels, maintaining CL homeostasis (30). Calpain may target proteins involved in this degradation pathway. Meanwhile, *Nr1h3* encodes LXRα, which facilitates reverse cholesterol transport (RCT) by promoting CL efflux via Abca1 and Abcg1 and by inducing bile acid synthesis via *Cyp7a1* (40-42). These results suggest that Capn4 may negatively regulate the RCT pathway via LXRα. Furthermore, LXRα nuclear translocation was enhanced in Capn4 knockdown livers, as shown by immunostaining. Because LXRα must localize to the nucleus to activate transcription of its target genes (32, 43), its nuclear translocation is essential for RCT induction. The nuclear import of LXRα has been reported to involve a basic amino acid–rich nuclear localization signal and a heterodimeric partner nuclear receptor, the latter of which is also known to be a substrate of Capn2 (32, 44). In addition, LXRα itself has been proposed as a calpain substrate in macrophages (45), further supporting the relationship between LXRα and calpain in this study.

Capn4 knockdown also led to increased expression of *Cpt1a*, the rate-limiting enzyme in mitochondrial β-oxidation, potentially counteracting ethanol-mediated suppression of fatty acid oxidation (Figure 4). Mitochondrial dysfunction is a key contributor to impaired β-oxidation in liver disease (46). This elevated serum FFA levels observed in Capn4 knockdown mice may reflect enhanced hepatic lipid export, potentially as a consequence of improved β-oxidation. In addition, the concurrent upregulation of *Dgat2* may represent a compensatory response aimed at mitigating lipotoxicity during transient FFA elevation (47). Moreover, ketone bodies are primarily produced in the liver from acetyl-CoA derived from β-oxidation and are subsequently transported to extrahepatic tissues for final oxidation (48). In non-alcoholic fatty liver disease, mitochondrial dysfunction is known to impair β-oxidation, leading to decreased serum β-hydroxybutyrate (βHB) levels (49). In this study, increased βHB levels in Capn4 knockdown mice is associated with improved mitochondrial β-oxidation. Therefore, Capn4 knockdown may contribute to improved mitochondrial function.

Early-stage ALD is characterized by the accumulation of LDs containing TG and cholesteryl esters (CE) (50, 51). There are two types of LDs: TG-rich and CE-rich, each with distinct protein profiles (52). Previous studies have demonstrated that LDs observed in microvesicular steatosis are predominantly enriched in CL rather than TG (29). Moreover, hepatic CL overload has been shown to induce microvesicular steatosis and concurrently impair mitochondrial function, highlighting the pathological link between CL accumulation and mitochondrial dysfunction in the liver (53).

In ALD, mitochondrial impairment contributes to excessive reactive oxygen species generation and endoplasmic reticulum stress (54). Ethanol-induced disruption of CL metabolism may further exacerbate these processes. This data support the notion that LXRα and HMGCR may be direct substrates of Capn4. As a mechanism underlying disease improvement, Capn4 knockdown reprograms hepatic homeostasis in response to ethanol, particularly in lipid metabolism centered on LXRα and HMGCR, improved mitochondrial function, and ultimately attenuation of microvesicular steatosis in ethanol-treated mice. Hepatic lipid metabolism involves multiple intersecting pathways, and the pathogenesis of ALD is highly complex.

Numerous calpain substrates remain unidentified. Elucidating calpain-regulated mechanisms involved in LXRα nuclear translocation and HMGCR degradation may provide mechanistic insights into ALD and inform therapeutic strategies. Selective inhibition of Capn4 may represent a novel therapeutic approach for restoring cholesterol homeostasis and preventing disease progression in ALD.

## Supporting information

graphical abstract

## Abbreviations

ASHI: alcohol-related steatohepatitis
Capns1I: Calpain small subunit 1
RTPCRI: reverse-transcriptase polymerase chain reaction
SDSI: sodium dodecyl sulfate
CysPCI: Calpain Cysteine Protease Core
Tnfα: Tumor necrosis factor-alpha
Pai-1I: Plasminogen activator inhibitor-1
FasnI: Fatty acid synthase
Ly6gI: Lymphocyte Antigen 6 Complex, Locus G
F4/80I: EGF-Like Module-Containing Mucin-Like Hormone Receptor-Like 1
BSAI: Bovine Serum Albumin
HMGCRI: 3-Hydroxy-3-Methylglutaryl-CoA Reductase
Dgat2I: Diacylglycerol o-acyltransferase 2
Cyp7a1I: Cytochrome p450 family 7 subfamily a member 1
LcatI: Lecithin-cholesterol acyltransferase
Srebf2I: Sterol regulatory element-binding factor 2
LXRαI: Liver X Receptor Alpha
ABCA1I: ATP-Binding Cassette Transporter A1
ABCG1I: ATP-Binding Cassette Transporter G1
SREBP2I: Sterol Regulatory Element-Binding Protein 2
LDLR I: Low-Density Lipoprotein Receptor
ROSI: Reactive Oxygen Species
ER I: Endoplasmic Reticulum
E11.5I: Embryonic day 11.5

## Disclosures

### Declaration of AI and AI-assisted technologies in the writing process

During the preparation of this work the authors used Claude (Anthropic) in order to assist with manuscript editing, grammar refinement, and structural organization of written content. After using this tool, the authors reviewed and edited the content as needed and take full responsibility for the content of the publication.

## Reference List

1. Staufer K, Stauber RE. Steatotic Liver Disease: Metabolic Dysfunction, Alcohol, or Both? Biomedicines 2023;11.

2. Seitz HK, Bataller R, Cortez-Pinto H, Gao B, Gual A, Lackner C, Mathurin P, et al. Alcoholic liver disease. Nat Rev Dis Primers 2018;4:16.

3. Parker R, Kim SJ, Gao B. Alcohol, adipose tissue and liver disease: mechanistic links and clinical considerations. Nat Rev Gastroenterol Hepatol 2018;15:50–59.

4. Stickel F, Hoehn B, Schuppan D, Seitz HK. Review article: Nutritional therapy in alcoholic liver disease. Aliment Pharmacol Ther 2003;18:357–373.

5. Bataller R, Arteel GE, Moreno C, Shah V. Alcohol-related liver disease: Time for action. J Hepatol 2019;70:221–222.

6. Sato T, Head KZ, Li J, Dolin CE, Wilkey D, Skirtich N, Smith K, et al. Fibrosis resolution in the mouse liver: Role of Mmp12 and potential role of calpain 1/2. Matrix Biol Plus 2023;17:100127.

7. Li J, Sato T, Hernandez-Tejero M, Beier JI, Sayed K, Benos PV, Wilkey DW, et al. The plasma degradome reflects later development of NASH fibrosis after liver transplant. Sci Rep 2023;13:9965.

8. Zhang M, Wang G, Peng T. Calpain-Mediated Mitochondrial Damage: An Emerging Mechanism Contributing to Cardiac Disease. Cells 2021;10.

9. Miyazaki T, Miyazaki A. Dysregulation of Calpain Proteolytic Systems Underlies Degenerative Vascular Disorders. J Atheroscler Thromb 2018;25:1–15.

10. Ge W, Hou C, Zhang W, Guo X, Gao P, Song X, Gao R, et al. Mep1a contributes to Ang II-induced cardiac remodeling by promoting cardiac hypertrophy, fibrosis and inflammation. J Mol Cell Cardiol 2021;152:52–68.

11. Letavernier E, Zafrani L, Perez J, Letavernier B, Haymann JP, Baud L. The role of calpains in myocardial remodelling and heart failure. Cardiovasc Res 2012;96:38–45.

12. Argemi J, Latasa MU, Atkinson SR, Blokhin IO, Massey V, Gue JP, Cabezas J, et al. Defective HNF4alpha-dependent gene expression as a driver of hepatocellular failure in alcoholic hepatitis. Nat Commun 2019;10:3126.

13. Bertola A, Mathews S, Ki SH, Wang H, Gao B. Mouse model of chronic and binge ethanol feeding (the NIAAA model). Nat Protoc 2013;8:627–637.

14. Massey VL, Dolin CE, Poole LG, Hudson SV, Siow DL, Brock GN, Merchant ML, et al. The hepatic “matrisome” responds dynamically to injury: Characterization of transitional changes to the extracellular matrix in mice. Hepatology 2017;65:969–982.

15. Isaacson RH, Beier JI, Khoo NK, Freeman BA, Freyberg Z, Arteel GE. Olanzapine-induced liver injury in mice: aggravation by high-fat diet and protection with sulforaphane. J Nutr Biochem 2020;81:108399.

16. Goll DE, Thompson VF, Li H, Wei W, Cong J. The calpain system. Physiol Rev 2003;83:731–801.

17. Ono Y, Sorimachi H. Calpains: an elaborate proteolytic system. Biochim Biophys Acta 2012;1824:224–236.

18. Miyazaki T, Taketomi Y, Saito Y, Hosono T, Lei XF, Kim-Kaneyama JR, Arata S, et al. Calpastatin counteracts pathological angiogenesis by inhibiting suppressor of cytokine signaling 3 degradation in vascular endothelial cells. Circ Res 2015;116:1170–1181.

19. Zhang C, Bai DS, Huang XY, Shi GM, Ke AW, Yang LX, Yang XR, et al. Prognostic significance of Capn4 overexpression in intrahepatic cholangiocarcinoma. PLoS One 2013;8:e54619.

20. Jang I, Menon S, Indra I, Basith R, Beningo KA. Calpain Small Subunit Mediated Secretion of Galectin-3 Regulates Traction Stress. Biomedicines 2024;12.

21. Bai DS, Dai Z, Zhou J, Liu YK, Qiu SJ, Tan CJ, Shi YH, et al. Capn4 overexpression underlies tumor invasion and metastasis after liver transplantation for hepatocellular carcinoma. Hepatology 2009;49:460–470.

22. Shields DC, Schaecher KE, Saido TC, Banik NL. A putative mechanism of demyelination in multiple sclerosis by a proteolytic enzyme, calpain. Proc Natl Acad Sci U S A 1999;96:11486–11491.

23. Ki SH, Park O, Zheng M, Morales-Ibanez O, Kolls JK, Bataller R, Gao B. Interleukin-22 treatment ameliorates alcoholic liver injury in a murine model of chronic-binge ethanol feeding: role of signal transducer and activator of transcription 3. Hepatology 2010;52:1291–1300.

24. Warner J, Hardesty J, Song Y, Sun R, Deng Z, Xu R, Yin X, et al. Fat-1 Transgenic Mice With Augmented n3-Polyunsaturated Fatty Acids Are Protected From Liver Injury Caused by Acute-On-Chronic Ethanol Administration. Front Pharmacol 2021;12:711590.

25. Hoebinger C, Rajcic D, Silva B, Hendrikx T. Chronic-binge ethanol feeding aggravates systemic dyslipidemia in Ldlr(-/-) mice, thereby accelerating hepatic fibrosis. Front Endocrinol (Lausanne) 2023;14:1148827.

26. Bertola A, Park O, Gao B. Chronic plus binge ethanol feeding synergistically induces neutrophil infiltration and liver injury in mice: a critical role for E-selectin. Hepatology 2013;58:1814–1823.

27. Alamri H, Patterson NH, Yang E, Zoroquiain P, Lazaris A, Chaurand P, Metrakos P. Mapping the triglyceride distribution in NAFLD human liver by MALDI imaging mass spectrometry reveals molecular differences in micro and macro steatosis. Anal Bioanal Chem 2019;411:885–894.

28. Fromenty B, Pessayre D. Impaired mitochondrial function in microvesicular steatosis. Effects of drugs, ethanol, hormones and cytokines. J Hepatol 1997;26 Suppl 2:43–53.

29. Minamikawa T, Ichimura-Shimizu M, Takanari H, Morimoto Y, Shiomi R, Tanioka H, Hase E, et al. Molecular imaging analysis of microvesicular and macrovesicular lipid droplets in non-alcoholic fatty liver disease by Raman microscopy. Sci Rep 2020;10:18548.

30. Shi Q, Chen J, Zou X, Tang X. Intracellular Cholesterol Synthesis and Transport. Front Cell Dev Biol 2022;10:819281.

31. Omkumar RV, Mehta PP, Kurup CK, Ramasarma T. Preparation of a soluble 58 kDa-3-hydroxy-3-methylglutaryl CoA reductase from liver microsomes and its inhibition by ethoxysilatrane, a hypocholesterolemic compound. Mol Cell Biochem 1992;110:145–153.

32. Prufer K, Boudreaux J. Nuclear localization of liver X receptor alpha and beta is differentially regulated. J Cell Biochem 2007;100:69–85.

33. You M, Arteel GE. Effect of ethanol on lipid metabolism. J Hepatol 2019;70:237–248.

34. Arthur JS, Elce JS, Hegadorn C, Williams K, Greer PA. Disruption of the murine calpain small subunit gene, Capn4: calpain is essential for embryonic development but not for cell growth and division. Mol Cell Biol 2000;20:4474–4481.

35. Schulman IG. Liver X receptors link lipid metabolism and inflammation. FEBS Lett 2017;591:2978–2991.

36. Goldstein JL, DeBose-Boyd RA, Brown MS. Protein sensors for membrane sterols. Cell 2006;124:35–46.

37. Marquart TJ, Allen RM, Ory DS, Baldan A. miR-33 links SREBP-2 induction to repression of sterol transporters. Proc Natl Acad Sci U S A 2010;107:12228–12232.

38. Madison BB. Srebp2: A master regulator of sterol and fatty acid synthesis. J Lipid Res 2016;57:333–335.

39. Zhang M, Ji J, Song J, An C, Pei W, Fan Q, Zuo L, et al. Current Therapeutic Targets for Alcohol-Associated Liver Disease. Am J Pathol 2025.

40. Wagner BL, Valledor AF, Shao G, Daige CL, Bischoff ED, Petrowski M, Jepsen K, et al. Promoter-specific roles for liver X receptor/corepressor complexes in the regulation of ABCA1 and SREBP1 gene expression. Mol Cell Biol 2003;23:5780–5789.

41. Peet DJ, Turley SD, Ma W, Janowski BA, Lobaccaro JM, Hammer RE, Mangelsdorf DJ. Cholesterol and bile acid metabolism are impaired in mice lacking the nuclear oxysterol receptor LXR alpha. Cell 1998;93:693–704.

42. Naik SU, Wang X, Da Silva JS, Jaye M, Macphee CH, Reilly MP, Billheimer JT, et al. Pharmacological activation of liver X receptors promotes reverse cholesterol transport in vivo. Circulation 2006;113:90–97.

43. Miller A, Crumbley C, Prufer K. The N-terminal nuclear localization sequences of liver X receptors alpha and beta bind to importin alpha and are essential for both nuclear import and transactivating functions. Int J Biochem Cell Biol 2009;41:834–843.

44. Matsushima-Nishiwaki R, Shidoji Y, Nishiwaki S, Moriwaki H, Muto Y. Limited degradation of retinoid X receptor by calpain. Biochem Biophys Res Commun 1996;225:946–951.

45. Miyazaki T, Miyazaki A. Emerging roles of calpain proteolytic systems in macrophage cholesterol handling. Cell Mol Life Sci 2017;74:3011–3021.

46. Wajner M, Amaral AU. Mitochondrial dysfunction in fatty acid oxidation disorders: insights from human and animal studies. Biosci Rep 2015;36:e00281.

47. Listenberger LL, Han X, Lewis SE, Cases S, Farese RV, Jr., Ory DS, Schaffer JE. Triglyceride accumulation protects against fatty acid-induced lipotoxicity. Proc Natl Acad Sci U S A 2003;100:3077–3082.

48. Puchalska P, Crawford PA. Multi-dimensional Roles of Ketone Bodies in Fuel Metabolism, Signaling, and Therapeutics. Cell Metab 2017;25:262–284.

49. Moore MP, Shryack G, Alessi I, Wieschhaus N, Meers GM, Johnson SA, Wheeler AA, et al. Relationship between serum beta-hydroxybutyrate and hepatic fatty acid oxidation in individuals with obesity and NAFLD. Am J Physiol Endocrinol Metab 2024;326:E493–E502.

50. Gao B, Bataller R. Alcoholic liver disease: pathogenesis and new therapeutic targets. Gastroenterology 2011;141:1572–1585.

51. Mashek DG, Khan SA, Sathyanarayan A, Ploeger JM, Franklin MP. Hepatic lipid droplet biology: Getting to the root of fatty liver. Hepatology 2015;62:964–967.

52. Khor VK, Ahrends R, Lin Y, Shen WJ, Adams CM, Roseman AN, Cortez Y, et al. The proteome of cholesteryl-ester-enriched versus triacylglycerol-enriched lipid droplets. PLoS One 2014;9:e105047.

53. Dominguez-Perez M, Simoni-Nieves A, Rosales P, Nuno-Lambarri N, Rosas-Lemus M, Souza V, Miranda RU, et al. Cholesterol burden in the liver induces mitochondrial dynamic changes and resistance to apoptosis. J Cell Physiol 2019;234:7213–7223.

54. Garcia-Ruiz C, Kaplowitz N, Fernandez-Checa JC. Role of Mitochondria in Alcoholic Liver Disease. Curr Pathobiol Rep 2013;1:159–168.

