## Supplementary document for "Calpain-4 Knockdown Modulates Cholesterol Metabolism and LXRα Nuclear Localization in Alcohol-Related Liver Disease"

[1]Department of Medicine, Division of Gastroenterology, Hepatology and Nutrition, [2]Pittsburgh Liver Research Center, University of Pittsburgh, [3]Department of Environmental and Occupational Health, [4]Pharmacology and Chemical Biology, University of Pittsburgh, [5]Organ Pathobiology and therapeutics institute, University of Pittsburgh, [6] Department of Internal Medicine, Liver Unit, Clinical University of Navarra, Navarra, Spain, [7]Institut d'Investigacions Biomediques August Pi i Sunyer (IDIBAPS), University of Barcelona, Barcelona, Spain.

Send all correspondence to: Gavin E. Arteel, PhD, FAASLD
Thomas E. Starzl Biomedical Science Tower
West 1144
200 Lothrop Street
Pittsburgh, PA 15213


### Supplementary Experimental Procedures

Information on rtPCR primers and probes, chemical assays, and antibodies used in this study are summarized in Supplementary Tables 1–3.

In addition, Supplementary Table 4 lists the top canonical pathways and upstream regulators identified by Ingenuity Pathway Analysis (IPA) for each of the three group comparisons (EtOH vs Control, Capn4KD vs Control, and EtOH+Capn4KD vs EtOH) from the RNA-seq dataset. Pathways are ranked by –log10 p-value, and z-scores are provided where available to indicate predicted activation or inhibition.

### Supplementary Results

Effect of ethanol exposure and Capn4 knockdown on indices of liver injury.

In addition to body weight and liver weight, we measured serum AST level, the expression of proinflammatory gene *(Tnfα*), and immune cell markers including *F4/80* (Kupffer cells and monocytes) and *Cd68* (inflammatory macrophages) to assess liver injury. Ethanol significantly reduced body weight gain, whereas no significant differences in the liver weight–to–body weight ratio were observed between groups. Although previous studies have reported increased serum AST level and *Tnfα* and *F4/80* expression following ethanol exposure (1, 2), these results did not show a significant change (Figure S1). In addition, Capn4 knockdown in ethanol-treated mice significantly increased expression of *F4/80* and *Cd68* (Figure S1). Previous studies have reported a link between macrophages and lipid metabolism, including β-oxidation (3, 4). These findings suggest that calpain may be involved in the relationship between macrophage activation and lipid accumulation.

Effect of ethanol exposure and Capn4 knockdown on Lipid Metabolism.

We examined genes involved in lipid metabolism, including *Fasn* and *Lcat*. Despite prior findings that ethanol promotes lipogenesis via upregulation of *Fasn*, there were no significant differences in this study. Capn4 knockdown in ethanol-treated mice did not alter the expression of these genes (Figure S2A). In addition, we analyzed proteins related to cholesterol metabolism, including VCP, ABCA1 and ABCG1. VCP is critical for the ER-associated degradation of HMG-CoA reductase and has been reported as a calpain substrate (5). In macrophage, there has been also reported that ABCA1 and ABCG1 are substrates of calpain (6). However, Capn4 knockdown in ethanol-treated mice did not alter the expression of these proteins.

Transcriptomic Analysis Reveals Reprogramming of Cholesterol Metabolism by Capn4 Knockdown Under Ethanol Stress.

To complement the primary findings presented in the main manuscript, we provide supplementary canonical pathway and upstream regulator analyses for the three group comparisons: EtOH vs Control, Capn4KD vs Control, and EtOH+Capn4KD vs EtOH. These analyses revealed broader transcriptomic alterations in addition to the cholesterol metabolism pathways emphasized in the main text. In the EtOH vs Control comparison, oxidative stress-related pathways—including NRF2-mediated Oxidative Stress Response (z-score: –1.41) and Acute Phase Response Signaling (z-score: –3.00)—were suppressed, in addition to the marked activation of biosynthetic programs such as Cholesterol Biosynthesis (z-score: 4.24). These findings are consistent with previous studies showing that ethanol exposure is closely associated with oxidative stress (7).

In the Capn4KD vs Control comparison, there was profound suppression of anabolic programs related to glucose and lipid metabolism (MLXIPL, z-score: –7.48), as well as protein synthesis pathways such as Response of EIF2AK4 (GCN2) to amino acid deficiency (z-score: –7.76) and Eukaryotic Translation Initiation (z-score: –8.25).

In contrast, the EtOH+Capn4KD vs EtOH comparison showed no significant changes in the anabolic responses observed in Capn4KD vs Control, but instead demonstrated suppression of Cholesterol biosynthetic pathways and upstream regulators previously activated by ethanol. These findings provide evidence that cholesterol metabolism undergoes reprogramming in response to ethanol exposure.

Reference List

1. Mathur M, Yeh YT, Arya RK, Jiang L, Pornour M, Chen W, Ma Y, et al. Adipose lipolysis is important for ethanol to induce fatty liver in the National Institute on Alcohol Abuse and Alcoholism murine model of chronic and binge ethanol feeding. Hepatology 2023;77:1688-1701.
2. Bertola A, Park O, Gao B. Chronic plus binge ethanol feeding synergistically induces neutrophil infiltration and liver injury in mice: a critical role for E-selectin. Hepatology 2013;58:1814-1823.
3. Yan J, Horng T. Lipid Metabolism in Regulation of Macrophage Functions. Trends Cell Biol 2020;30:979-989.
4. Huang SC, Everts B, Ivanova Y, O'Sullivan D, Nascimento M, Smith AM, Beatty W, et al. Cell-intrinsic lysosomal lipolysis is essential for alternative activation of macrophages. Nat Immunol 2014;15:846-855.
5. Matsushima-Nishiwaki R, Shidoji Y, Nishiwaki S, Moriwaki H, Muto Y. Limited degradation of retinoid X receptor by calpain. Biochem Biophys Res Commun 1996;225:946-951.
6. Miyazaki T, Miyazaki A. Emerging roles of calpain proteolytic systems in macrophage cholesterol handling. Cell Mol Life Sci 2017;74:3011-3021.
7. Albano E. Alcohol, oxidative stress and free radical damage. Proc Nutr Soc 2006;65:278-290.

### Figure legends

**Supplementary Figure S1. Effect of ethanol exposure and Capn4 knockdown on indices of liver injury.**

To assess liver injury, we measured serum AST level, the expression of proinflammatory gene, *Tnfα*, and immune cell markers, *Cd68* (inflammatory macrophages), and *F4/80* (Kupffer cells and monocytes) in addition to body weight and liver weight. PF, pair-fed; EtOH, ethanol; AST, aspartate aminotransferase.

**Supplementary Figure S2. Effect of ethanol exposure and Capn4 knockdown on lipid metabolism.**

Panel A: The expression of key lipid metabolism genes was determined. These included genes involved in fatty acid metabolism (*Fasn*) and cholesterol homeostasis (*Lcat*). Panel B: The protein levels of mediators of cholesterol metabolism were determined. Specifically, the expression of ABCA1, ABCG1 and VCP was determined.

**Supplementary Figure S3. Canonical pathways and upstream regulators identified by Ingenuity Pathway Analysis (IPA) in each group comparison from the RNA-seq dataset.**

(A) EtOH vs Control, (B) Capn4KD vs Control, and (C) EtOH+Capn4KD vs EtOH. Pathways are ranked by –log10 p-value, and z-scores are shown where available. Positive z-scores represent predicted activation, whereas negative z-scores represent predicted inhibition. NA indicates no z-score available. IPA, Ingenuity Pathway Analysis; KD, knockdown.

**Supplementary Table 4. Enriched canonical pathways and upstream regulators (IPA analysis).**

Canonical pathways and upstream regulators identified by Ingenuity Pathway Analysis (IPA) in each group comparison from the RNA-seq dataset. Pathways are ranked by –log10 p-value. Predicted activation or inhibition is indicated by z-score, with positive values representing activation and negative values representing inhibition. NA indicates that a z-score could not be calculated. Gene symbols are displayed in uppercase (e.g., SREBF2, ABCA1) based on IPA output, while gene symbols in the main text follow mouse nomenclature (e.g., *Srebf2, Abca1*).
