## Supplementary Table S1 for "Calpain-4 Knockdown Modulates Cholesterol Metabolism and LXRα Nuclear Localization in Alcohol-Related Liver Disease"

**Supplemental Table 1-Product information for primers used in RT-PCR**

| **Gene name** | **Supplier** | **Cat. No.** |
| --- | --- | --- |
| *Capn1* | Thermofisher | Mm00482964_m1 |
| *Capn2* | Thermofisher | Mm00486669_m1 |
| *CapnS1* | Thermofisher | Mm00501568_m1 |
| *Cast* | Thermofisher | Mm01345276_Mh |
| *Tnfa* | Thermofisher | Mm00443258_m1 |
| *Serpine* | Thermofisher | Rn01481341_m1 |
| *Ly6g* | Thermofisher | Mm04934123_m1 |
| *Adgre1* | Thermofisher | Mn00802529_m1 |
| *Cd68* | Thermofisher | Mm03047343_m1 |
| *Srebf2* | Thermofisher | Mm01306292_m1 |
| *Fasn* | Thermofisher | Mm00662319_m1 |
| *Cpt1a* | Thermofisher | Mm00550438_m1 |
| *Dgat2* | Thermofisher | Mm00499536_m1 |
| *Lcat* | Thermofisher | Mm01178820_m1 |
| *Cyp7a1* | Thermofisher | Mm00484150_m1 |
| *Nr1h3* | Thermofisher | Mm00443451_m1 |
| *Abca1* | Thermofisher | Mm01178820_m1 |
| *Abcg1* | Thermofisher | Mm00437390_m1 |
