## Supplementary Table S2 for "Calpain-4 Knockdown Modulates Cholesterol Metabolism and LXRα Nuclear Localization in Alcohol-Related Liver Disease"

**Supplemental Table 2-** **Drugs and Chemical assays used in this study**

| **Category** | **Name** | **Supplier** | **Cat No.** |
| --- | --- | --- | --- |
| Chemical assays | AST assay kit | Thermofisher | TR70121 |
|  | ALT assay kit | Thermofisher | TR71121 |
|  | Infinity™ Triglycerides Reagent | Thermofisher | TR22421 |
|  | Infinity™ Cholesterol Reagent | Thermofisher | TR13421 |
|  | Free Fatty Acid Assay Kit | Cell Biolabs, Inc. | STA-618 |
|  | Beta-Hydroxybutyrate Assay Kit (Colorimetric) | Abcam | ab83390 |
