## Supplementary Table S3 for "Calpain-4 Knockdown Modulates Cholesterol Metabolism and LXRα Nuclear Localization in Alcohol-Related Liver Disease"

**Supplemental Table 3-** **Primary antibodies used in this study**

| **Antibody** | **Supplier** | **Cat No.** |
| --- | --- | --- |
| GAPDH | CELL SIGNALING | 5174S |
| Capns1 | LSBio | LS-C482635-30 |
| LXRα | Abcam | ab176323 |
| ABCA1 | Abcam | ab18180 |
| ABCG1 | Abcam | ab52617 |
| HMGCR | Invitrogen | XL3767433A |
| VCP | Proteintech | 10736-1-AP |
