## Supplementary Table S4 for "Calpain-4 Knockdown Modulates Cholesterol Metabolism and LXRα Nuclear Localization in Alcohol-Related Liver Disease"

**Supplemental Table 4-** **canonical pathways and upstream regulators (IPA analysis).**

**EtOH vs control -** **canonical pathways**

| Ingenuity Canonical Pathways | -log(p-value) | z-score | gene count |
| --- | --- | --- | --- |
| Cholesterol biosynthesis | 13.2 | 4.243 | 18 |
| Activation of gene expression by SREBF (SREBP) | 11.9 | 4.025 | 21 |
| Superpathway of Cholesterol Biosynthesis | 11.7 | 4.123 | 18 |
| LPS/IL-1 Mediated Inhibition of RXR Function | 11.2 | -2.065 | 62 |
| NRF2-mediated Oxidative Stress Response | 11 | -1.414 | 58 |
| Pulmonary Fibrosis Idiopathic Signaling Pathway | 10.4 | -1.664 | 64 |
| Regulation of lipid metabolism by PPARalpha | 9.57 | -0.522 | 33 |
| Molecular Mechanisms of Cancer | 9.09 | -0.271 | 123 |
| Phase I - Functionalization of compounds | 8.91 | 0.365 | 31 |
| Thrombin Signaling | 8.83 | -0.649 | 47 |
| Role of BRCA1 in DNA Damage Response | 8.59 | 0.728 | 25 |
| RHO GTPase cycle | 8.57 | 0.115 | 75 |
| Cholesterol Biosynthesis I | 8.52 | 3.162 | 10 |
| Cholesterol Biosynthesis II (via 24,25-dihydrolanosterol) | 8.52 | 3.162 | 10 |
| Cholesterol Biosynthesis III (via Desmosterol) | 8.52 | 3.162 | 10 |
| Cardiac Hypertrophy Signaling (Enhanced) | 8.51 | -1.99 | 85 |
| LXR/RXR Activation | 8.5 | 1.789 | 33 |
| Hepatic Fibrosis Signaling Pathway | 8.22 | -1.807 | 70 |
| Aryl Hydrocarbon Receptor Signaling | 8.09 | 1.414 | 40 |
| RAR Activation | 7.49 | -0.611 | 70 |
| Glycation Signaling Pathway | 7.29 | -1.372 | 44 |
| Protein Kinase A Signaling | 6.91 | 1.483 | 64 |
| Neutrophil degranulation | 6.8 | -1.053 | 73 |
| Role of Osteoclasts in Rheumatoid Arthritis Signaling Pathway | 6.67 | -1.443 | 54 |
| Acute Phase Response Signaling | 6.59 | -3 | 38 |
| Extracellular matrix organization | 6.44 | 1.177 | 26 |
| Chronic Myeloid Leukemia Signaling | 6.43 | -0.866 | 49 |
| GP6 Signaling Pathway | 6.4 | 1.89 | 29 |
| FXR/RXR Activation | 6.36 | 0.354 | 38 |
| Response of EIF2AK1 (HRI) to heme deficiency | 6.28 | -3 | 9 |

**Capn4 KD vs control -** **canonical pathways**

| Ingenuity Canonical Pathways | -log(p-value) | z-score | gene count |
| --- | --- | --- | --- |
| Response of EIF2AK4 (GCN2) to amino acid deficiency | 55.6 | -7.761 | 69 |
| Nonsense-Mediated Decay (NMD) | 50 | -7.761 | 69 |
| Eukaryotic Translation Elongation | 47 | -8.185 | 68 |
| Eukaryotic Translation Initiation | 44.9 | -8.25 | 73 |
| Eukaryotic Translation Termination | 44.5 | -8.062 | 66 |
| Selenoamino acid metabolism | 42.5 | -8.066 | 70 |
| Major pathway of rRNA processing in the nucleolus and cytosol | 41 | -8.488 | 77 |
| SRP-dependent cotranslational protein targeting to membrane | 39.1 | -8.062 | 66 |
| Signaling by ROBO receptors | 36.2 | -1.886 | 83 |
| EIF2 Signaling | 35.7 | -4.802 | 81 |
| Ribosomal Quality Control Signaling Pathway | 35.3 | -8.51 | 85 |
| Respiratory electron transport | 29.1 | -6.708 | 45 |
| Oxidative Phosphorylation | 22.3 | -6.164 | 43 |
| Mitochondrial translation | 21.5 | -6.325 | 40 |
| Sirtuin Signaling Pathway | 18.4 | 3.286 | 67 |
| Mitochondrial Dysfunction | 18.3 | 5.416 | 73 |
| Complex I biogenesis | 17.3 | -5.477 | 30 |
| Regulation of eIF4 and p70S6K Signaling | 16.3 | 0.378 | 49 |
| Granzyme A Signaling | 14.9 | 4.6 | 29 |
| mTOR Signaling | 14.6 | -0.577 | 50 |
| Estrogen Receptor Signaling | 13.1 | 2.777 | 71 |
| Cytoprotection by HMOX1 | 11.3 | -2.828 | 22 |
| Mitochondrial protein import | 10.1 | -4.69 | 22 |
| Complex IV assembly | 9.23 | -4.243 | 18 |
| Coronavirus Pathogenesis Pathway | 8.85 | 3.333 | 40 |
| TP53 Regulates Metabolic Genes | 8.38 | -3.128 | 23 |
| Hematoma Resolution Signaling Pathway | 7.65 | -4.218 | 43 |
| Parkinson's Signaling Pathway | 7.2 | 6.332 | 48 |
| Regulation of mitotic cell cycle | 6.68 | -2.837 | 21 |
| Regulation of mRNA stability by proteins that bind AU-rich elements | 6.59 | -3.273 | 21 |

**EtOH+Capn4 KD vs EtOH –** **canonical pathways**

| Ingenuity Canonical Pathways | -log(p-value) | z-score | gene count |
| --- | --- | --- | --- |
| HEY1 Signaling Pathway | 8.38 | 1.508 | 11 |
| Molecular Mechanisms of Cancer | 4.72 | 1.5 | 17 |
| Cholesterol biosynthesis | 4.7 | -2 | 4 |
| CDX Gastrointestinal Cancer Signaling Pathway | 4.56 | -1.414 | 8 |
| Superpathway of Cholesterol Biosynthesis | 4.45 | -2 | 4 |
| G alpha (i) signalling events | 3.75 | 2.121 | 8 |
| Pulmonary Healing Signaling Pathway | 3.65 | 0.378 | 7 |
| Sheddase Signaling Pathway | 3.61 | 1.89 | 7 |
| S100 Family Signaling Pathway | 3.54 | 1.941 | 14 |
| Factors Promoting Cardiogenesis in Vertebrates | 3.5 | 0 | 6 |
| NCAM signaling for neurite out-growth | 3.24 | 1 | 4 |
| Role of Osteoblasts in Rheumatoid Arthritis Signaling Pathway | 3.18 | 1.134 | 7 |
| Role of Osteoclasts in Rheumatoid Arthritis Signaling Pathway | 3.16 | -1.342 | 8 |
| Zn Homeostasis Signaling Pathway | 3.04 | 0.5 | 16 |
| Cellular Effects of Sildenafil (Viagra) | 3.02 | 0.302 | 12 |
| RAR Activation | 2.88 | 0.333 | 9 |
| Cerebral Malformation Signaling Pathway | 2.83 | 1.342 | 5 |
| Endocannabinoid Cancer Inhibition Pathway | 2.7 | -2 | 5 |
| Activation of gene expression by SREBF (SREBP) | 2.68 | N/A | 3 |
| Tuberculosis Latent Signaling Pathway | 2.66 | 0 | 4 |
| BMP signaling pathway | 2.66 | 0 | 4 |
| Transcriptional Regulatory Network in Embryonic Stem Cells | 2.49 | -0.447 | 5 |
| Glutaminergic Receptor Signaling Pathway (Enhanced) | 2.43 | 1.89 | 7 |
| Wound Healing Signaling Pathway | 2.39 | 0 | 6 |
| Post-translational protein phosphorylation | 2.39 | -2 | 4 |
| Class A/1 (Rhodopsin-like receptors) | 2.38 | 0.378 | 7 |
| CREB Signaling in Neurons | 2.37 | 1 | 10 |
| Tumor Microenvironment Pathway | 2.32 | 0.447 | 5 |
| p38 MAPK Signaling | 2.17 | 0 | 4 |
| Regulation of Insulin-like Growth Factor (IGF) transport and uptake by IGFBPs | 2.17 | -2 | 4 |

**EtOH vs control - upstream regulators**

| Upstream Regulator | Molecule Type | z-score | p-value of overlap |
| --- | --- | --- | --- |
| PPARA | ligand-dependent nuclear receptor | 0.048 | 1.03E-46 |
| AHR | ligand-dependent nuclear receptor | -1.2 | 5.64E-42 |
| TGFB1 | growth factor | -3.227 | 1.43E-37 |
| STAT5B | transcription regulator | 3.412 | 3.74E-36 |
| NFE2L2 | transcription regulator | -3.797 | 1.98E-35 |
| RORA | ligand-dependent nuclear receptor | 0.217 | 7.86E-31 |
| PPARG | ligand-dependent nuclear receptor | 2.017 | 1.46E-30 |
| SREBF2 | transcription regulator | 4.575 | 9.86E-30 |
| AGT | growth factor | -1.223 | 2.47E-29 |
| MYC | transcription regulator | -2.963 | 1.67E-28 |
| NR1I3 | ligand-dependent nuclear receptor | -4.699 | 5.88E-28 |
| TP63 | transcription regulator | -2.303 | 6.09E-27 |
| TP53 | transcription regulator | -2.689 | 6.19E-26 |
| CEBPB | transcription regulator | -0.063 | 9.32E-25 |
| FOXO1 | transcription regulator | -1.287 | 2.72E-24 |
| ESR2 | ligand-dependent nuclear receptor | 0.094 | 5.24E-24 |
| NR3C1 | ligand-dependent nuclear receptor | -1.453 | 5.04E-23 |
| IGF1 | growth factor | -1.006 | 1.1E-22 |
| PPARD | ligand-dependent nuclear receptor | 1.489 | 1.86E-22 |
| RORC | ligand-dependent nuclear receptor | 1 | 2.52E-22 |
| FOXO3 | transcription regulator | -2.451 | 3.49E-22 |
| CEBPA | transcription regulator | -2.221 | 3.66E-22 |
| NFAT5 | transcription regulator | -0.236 | 1.44E-21 |
| HUWE1 | transcription regulator | 2.665 | 2.62E-21 |
| RXRA | ligand-dependent nuclear receptor | 1.819 | 1.84E-20 |
| HNF4A | transcription regulator | 1.997 | 4.04E-20 |
| HMG20A | transcription regulator | 1.417 | 5.81E-20 |
| SREBF1 | transcription regulator | 3.233 | 6.06E-20 |
| HTT | transcription regulator | -1.801 | 9.32E-20 |
| SIRT1 | transcription regulator | -2.819 | 1.09E-19 |

**Capn4 KD vs control – upstream regulators**

| Upstream Regulator | Molecule Type | z-score | p-value of overlap |
| --- | --- | --- | --- |
| MLXIPL | transcription regulator | -7.842 | 3.63E-49 |
| LARP1 | translation regulator | 7.547 | 1.9E-46 |
| SPEN | transcription regulator | -6.856 | 4.83E-42 |
| CTNNB1 | transcription regulator | 2.183 | 3.13E-30 |
| MYC | transcription regulator | -4.941 | 2.27E-26 |
| FMR1 | translation regulator | 6.378 | 5.4E-26 |
| YAP1 | transcription regulator | 1.031 | 3.12E-24 |
| HNF4A | transcription regulator | 0.987 | 1.21E-23 |
| EIF6 | translation regulator | 5.555 | 3.62E-21 |
| TEAD1 | transcription regulator | -4.922 | 9.01E-17 |
| MYCN | transcription regulator | -5.653 | 1.68E-16 |
| ZHX2 | transcription regulator | 4.591 | 1.27E-11 |
| HIF1A | transcription regulator | 1.229 | 1.28E-11 |
| PPARA | ligand-dependent nuclear receptor | -0.926 | 5.9E-11 |
| RB1 | transcription regulator | -4.816 | 7.9E-11 |
| CLPB | transcription regulator | 3.051 | 1.29E-08 |
| RXRA | ligand-dependent nuclear receptor | 1.359 | 0.000000235 |
| IGF2BP1 | translation regulator | 1.231 | 0.000000316 |
| COPS5 | transcription regulator | 3.06 | 0.00000042 |
| AGT | growth factor | 0.897 | 0.000000427 |
| HUWE1 | transcription regulator | 1.187 | 0.000000437 |
| STAT5B | transcription regulator | 1.523 | 0.000000467 |
| NR3C1 | ligand-dependent nuclear receptor | -1.008 | 0.00000075 |
| PPARGC1A | transcription regulator | -1.232 | 0.000000923 |
| NR1I3 | ligand-dependent nuclear receptor | -1.165 | 0.00000197 |
| TP53 | transcription regulator | -0.245 | 0.00000244 |
| NRF1 | transcription regulator | -3.074 | 0.00000279 |
| HTT | transcription regulator | -0.327 | 0.0000029 |
| PITX2 | transcription regulator | -2.855 | 0.00000297 |
| MYCL | transcription regulator | -2.926 | 0.00000324 |

**EtOH + Capn4 KD vs EtOH – upstream regulators**

| Upstream Regulator | Molecule Type | z-score | p-value of overlap |
| --- | --- | --- | --- |
| TGFB1 | growth factor | 0.831 | 1.1E-10 |
| NPM1 | transcription regulator | 0.333 | 0.000000214 |
| SMARCB1 | transcription regulator | -1.134 | 0.000000305 |
| ARID1A | transcription regulator | -2.401 | 0.00000729 |
| SREBF2 | transcription regulator | -2.607 | 0.0000108 |
| PGR | ligand-dependent nuclear receptor | -1.947 | 0.0000185 |
| SREBF1 | transcription regulator | -0.918 | 0.0000276 |
| EGF | growth factor | 1.52 | 0.0000292 |
| FOS | transcription regulator | 0.816 | 0.000043 |
| KMT2D | transcription regulator | -0.552 | 0.000068 |
| AHR | ligand-dependent nuclear receptor | 0.214 | 0.000177 |
| TP63 | transcription regulator | 2.09 | 0.000193 |
| AGT | growth factor | -0.777 | 0.00022 |
| TGFB3 | growth factor | 0.816 | 0.000234 |
| NCOA2 | transcription regulator | -0.816 | 0.000327 |
| ARID2 | transcription regulator | 0 | 0.000359 |
| BMP10 | growth factor | 0.447 | 0.000387 |
| ESR2 | ligand-dependent nuclear receptor | -1.667 | 0.000407 |
| GDF2 | growth factor | 0 | 0.000478 |
| SMAD3 | transcription regulator | -0.053 | 0.000491 |
| BMP2 | growth factor | 2.149 | 0.000613 |
| STAT5B | transcription regulator | 2.121 | 0.000649 |
| CTNNB1 | transcription regulator | -0.513 | 0.000727 |
| IGF1 | growth factor | 0.733 | 0.000802 |
| QKI | transcription regulator | -1.253 | 0.000854 |
| TARDBP | transcription regulator | -2 | 0.000913 |
| FOXO4 | transcription regulator | 1.06 | 0.000961 |
| NFKB2 | transcription regulator | -0.962 | 0.000999 |
| ESRRA | transcription regulator | -1 | 0.00139 |
| STAT3 | transcription regulator | -0.047 | 0.00165 |
