## Supplementary figures and images for "Calpain-4 Knockdown Modulates Cholesterol Metabolism and LXRα Nuclear Localization in Alcohol-Related Liver Disease"

### Supplementary Figure S1

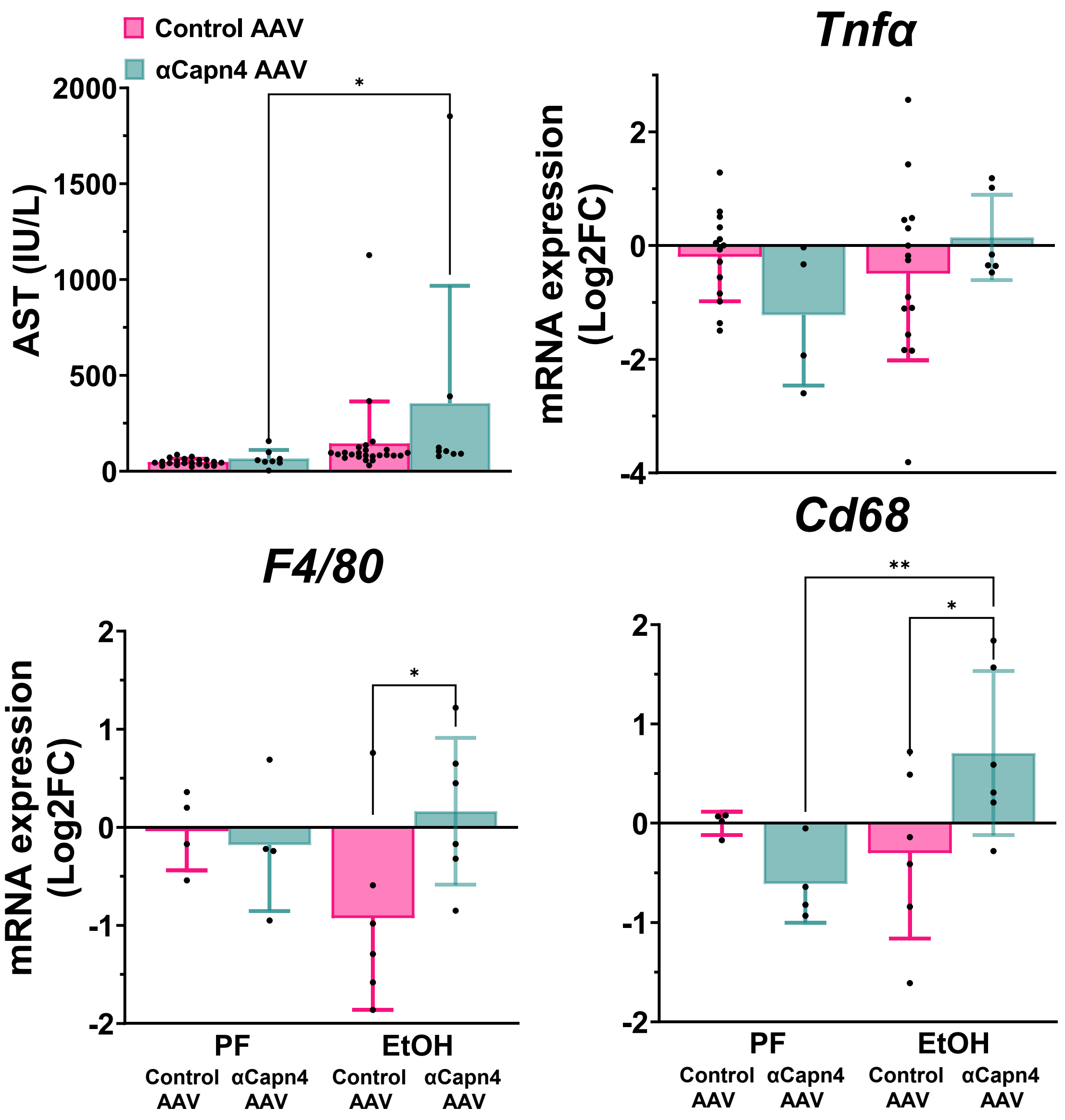

### Supplementary Figure S2

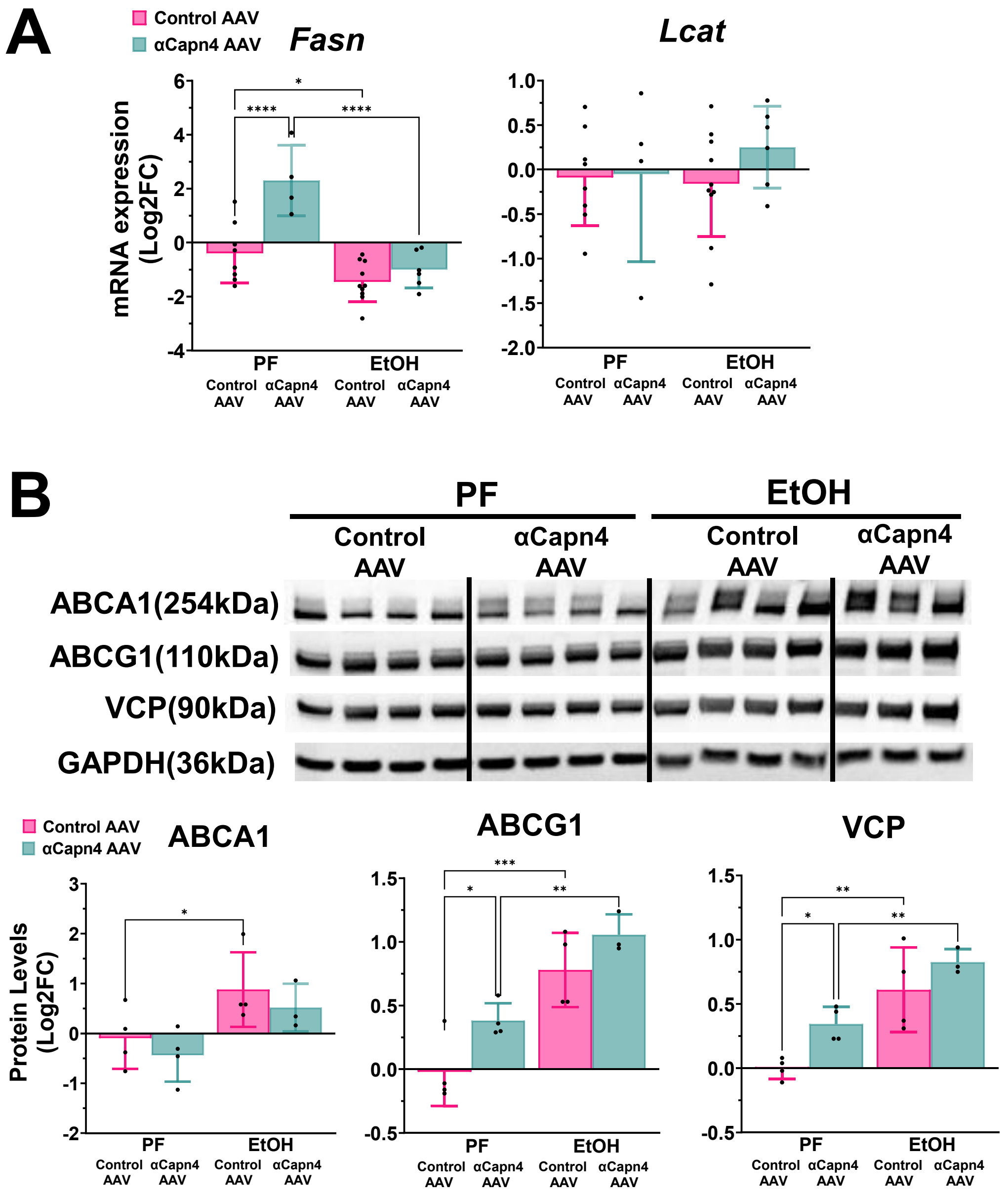

### Supplementary Figure S3

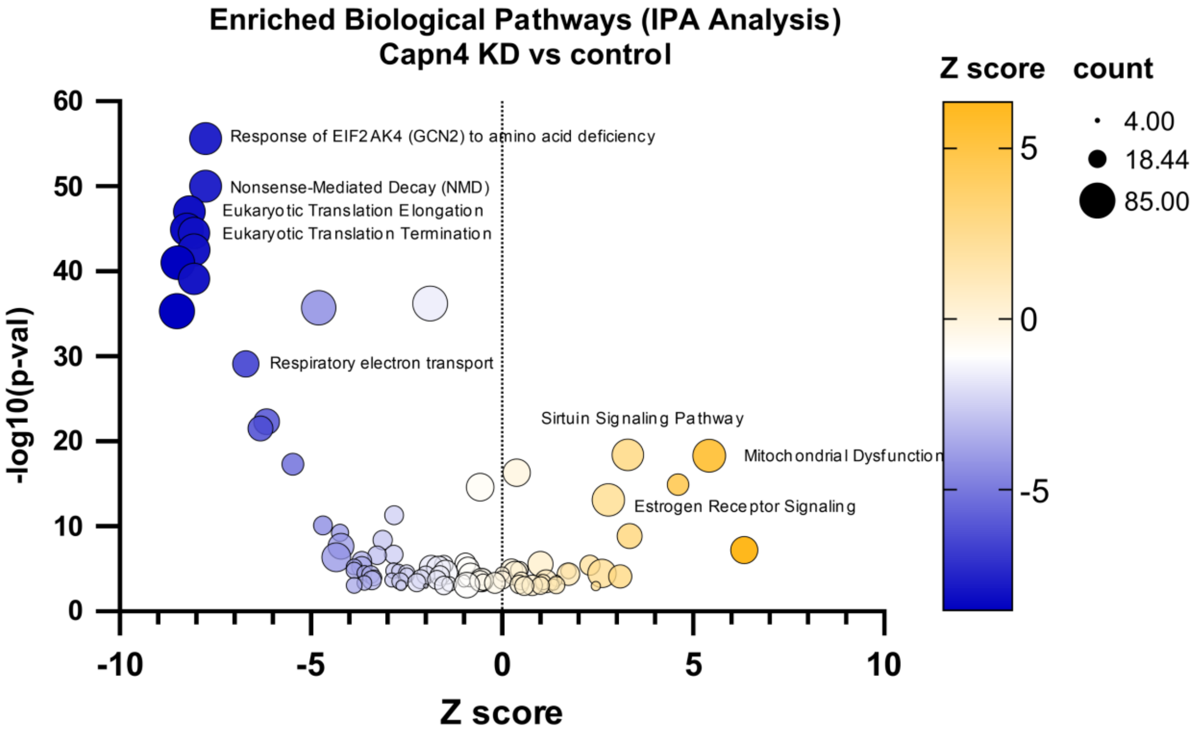
