## Supplementary material for "Calpain-4 Knockdown Modulates Cholesterol Metabolism and LXRα Nuclear Localization in Alcohol-Related Liver Disease": graphical abstract

### Calpain-4 Knockdown in ALD: Key Results

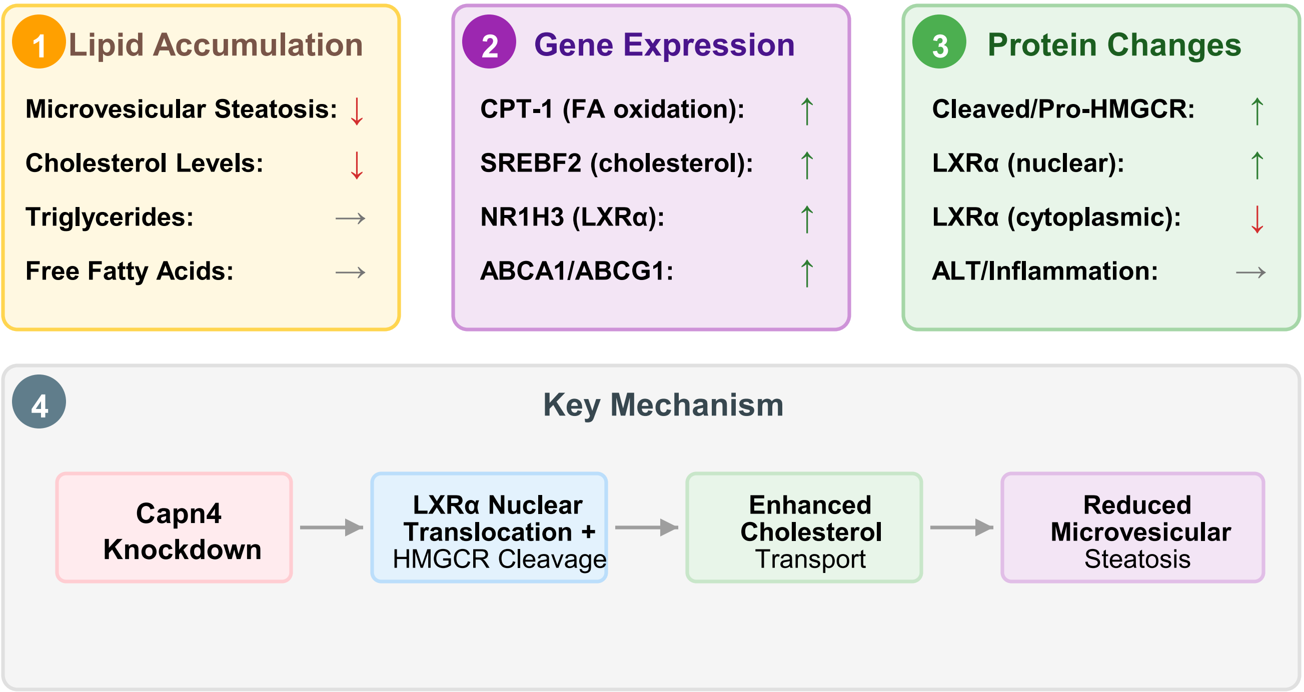

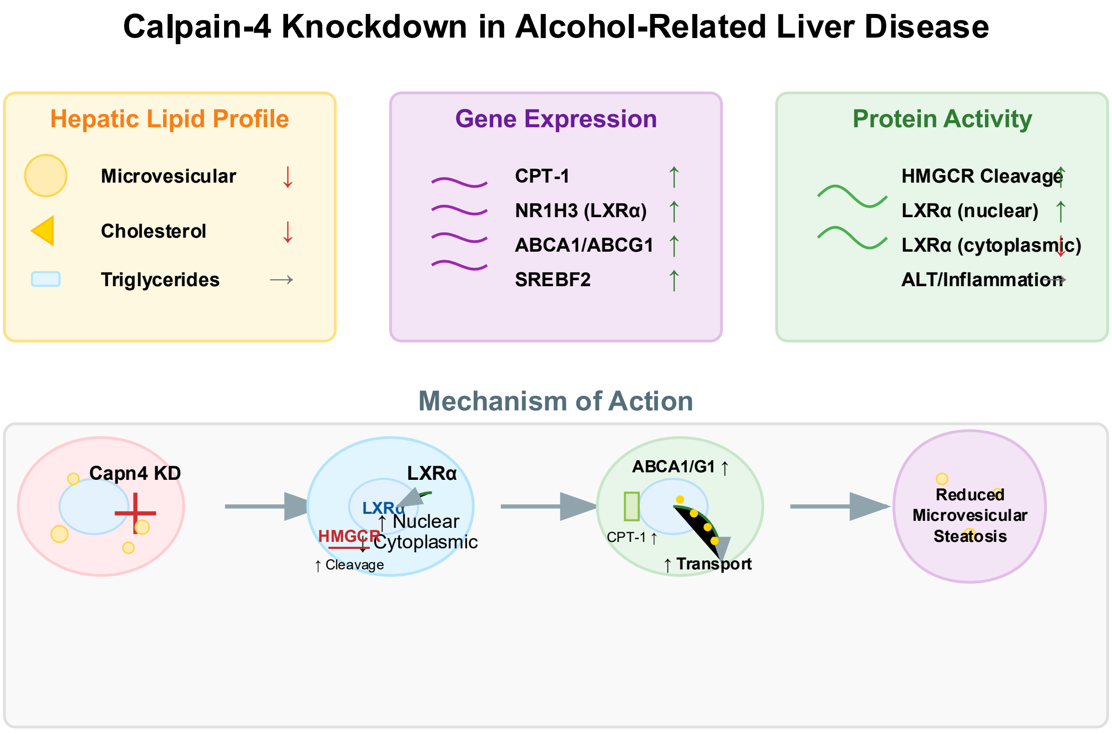


Kitano et al. "Calpain-4 Knockdown Modulates Cholesterol Metabolism and LXRα Nuclear Localization in Alcohol-Related Liver Disease"
